# A photoswitchable GPCR-based opsin for presynaptic silencing

**DOI:** 10.1101/2021.02.19.432008

**Authors:** Bryan A. Copits, Patrick R. O’Neill, Raaj Gowrishankar, Judy J. Yoo, Xenia Meshik, Kyle E. Parker, Skylar M. Spangler, Alexis M. Vasquez, Abigail J. Elerding, M. Christine Stander, Vani Kalyanaraman, Sherri K. Vogt, Vijay K. Samineni, N. Gautam, Roger K. Sunahara, Robert W. Gereau, Michael R. Bruchas

**Affiliations:** Washington University Pain Center, Washington University School of Medicine; St. Louis, MO; Department of Anesthesiology, Washington University School of Medicine; St. Louis, MO; Department of Neuroscience, Washington University School of Medicine; St. Louis, MO; Department of Pharmacology, University of California San Diego; San Diego, CA; Center of Excellence in the Neurobiology of Addiction, Pain, and Emotion; Departments of Anesthesiology and Pharmacology, University of Washington; Seattle, WA

## Abstract

Optical manipulations of genetically defined cell types have generated significant insights into the dynamics of neural circuits. While optogenetic activation has been relatively straightforward, rapid and reversible synaptic inhibition has been far more difficult to achieve. Instead of relying on unpredictable ion manipulations or slow photoactivatable toxins at axon terminals, we took a different approach to leverage the natural ability of inhibitory presynaptic GPCRs to silence synaptic transmission. Here we characterize parapinopsin (PPO), a photoswitchable non-visual opsin from lamprey pineal gland that couples to G_i/o_-signaling cascades. PPO can be rapidly activated by pulsed blue light, switched off with amber light, and is effective for repeated or prolonged inhibition. We developed viral vectors for cell-specific expression of PPO, which traffics very effectively in numerous neuron types. At presynaptic terminals, PPO can silence glutamate release and suppress dopamine-dependent reward and cocaine place preference behaviors *in vivo*. PPO immediately fills a significant gap in the neuroscience toolkit for rapid and reversible synaptic inhibition, and has broader utility for achieving spatiotemporal control of inhibitory GPCR signaling cascades in other biological and pharmacological applications.

## INTRODUCTION

The development of diverse molecular tools to monitor and control defined cell types has greatly accelerated our understanding of fundamental biological processes. Within the field of neuroscience, these tools have granted access to the circuit dynamics underlying complex behaviors. Approaches to manipulate neuronal activity have largely focused on optogenetic strategies using light-activated ion channels and pumps (Copits et al., 2016; Kim et al., 2017; Rost et al., 2017; Wiegert et al., 2017), or chemogenetic implementations with engineered G protein coupled receptors (GPCRs) (Atasoy and Sternson, 2018; Burnett and Krashes, 2016; Roth, 2016). The primary strength of optogenetic activators has been their high degree of spatiotemporal precision in controlling neuronal firing, and more importantly their distinct axonal projections, at millisecond timescales. However, optogenetic inhibition using ion-based opsins at synaptic terminals has proven to be far more problematic. These challenges are largely due to biophysical limitations that can cause direct depolarization or rebound spiking (Mahn et al., 2016, 2018; Messier et al., 2018; Raimondo et al., 2012; Wiegert et al., 2017). An alternative strategy for inhibiting synaptic projections used chemogenetic receptors to take advantage of the natural ability of G_i_-coupled GPCRs to silence synaptic transmission (Stachniak et al., 2014). However, these chemogenetic tools lack spatial and temporal precision compared to optical approaches.

Engineered rhodopsin-GPCR (opto-XR) chimeras, in which light sensitivity is granted to different GPCRs by splicing them to the light-sensitive visual protein rhodopsin, bridges opto- and chemogenetic approaches to gain spatiotemporal optical control of physiological signaling cascades (Airan et al., 2009; Kleinlogel, 2016). However, existing opto-XRs are limited by the extremely high photosensitivity and irreversible activation of the rhodopsin chromophore (Ernst et al., 2014). Evolution has generated tremendous diversity in visual and non-visual opsins throughout the animal kingdom (Davies et al., 2010; Koyanagi and Terakita, 2014; Marshall et al., 2015; Terakita et al., 2015). We reasoned that some of these opsins may have evolved unique spectral and/or kinetic properties that could make them superior alternative candidates to chemogenetic and rhodopsin-based optogenetic approaches for engaging endogenous signaling cascades to manipulate neural circuits. The ideal opto-XR would be (1) rapidly reversible, (2) exhibit moderate light sensitivity to avoid unwanted opsin activation by ambient light, and (3) possess biocompatible spectral properties that overlap with existing optogenetic light sources. Additionally, opsin coupling to inhibitory GPCR signaling cascades that have been evolutionarily co-opted to suppress neurotransmitter release, would directly address a major limitation with currently available tools – silencing synaptic projections (Wiegert et al., 2017).

We searched and initially tested various opsins that might possess these features, and identified lamprey parapinopsin (PPO) as a potential candidate optogenetic tool (Kawano-Yamashita et al., 2015; Koyanagi and Terakita, 2014; Koyanagi et al., 2004, 2017; O’Neill et al., 2018). PPO is a non-visual G protein coupled opsin expressed in the lamprey pineal gland that exhibits bistable photoswitching between on and off states when irradiated with ultra-violet (UV) *vs.* green/amber light (Kawano-Yamashita et al., 2015; Koyanagi et al., 2004). This bistable optical sensitivity is thought to confer “color discrimination” abilities to sense wavelength changes during diurnal or seasonal changes in these ancient vertebrates (Dodt and Meissl, 1982; Koyanagi et al., 2004; Morita et al., 1992; Uchida and Morita, 1994). We reasoned that this unique photoswitchable property might be harnessed for more precise spatiotemporal modulation of neuronal circuits.

Here we establish PPO as a novel photoswitchable GPCR-based opsin for rapid and reversible optical control of inhibitory G protein signaling pathways and thus synaptic silencing in neural circuits. We show that pulsed blue light stimulation of PPO activates G protein cascades at cellular and subcellular resolution and can be rapidly switched off with amber light. PPO can be activated by multiphoton stimulation and multiplexed with red-shifted indicators of neuronal activity. PPO couples to neuronal G proteins to inhibit voltage-gated calcium channels in an equivalent manner to native G_i_-coupled GABA_B_ receptors. We found that PPO rapidly and reliably couples to inhibitory signaling pathways for either repeated or prolonged optical modes of engagement. We developed Cre-dependent viral vectors for cell-specific expression in the brain, and found that PPO traffics very efficiently in multiple neuron types to both dendrites and presynaptic terminals, where it can be effectively stimulated at the latter site to silence synaptic transmission. Finally, we go on to demonstrate that, when illuminated at terminal sites, PPO can suppress dopamine-dependent reward seeking behaviors and cocaine-induced preference mediated by ventral tegmental area dopamine neuron projections to the nucleus accumbens. Thus, PPO represents a unique GPCR-based tool that bridges the gap between optogenetics and chemogenetics and allows for projection-specific presynaptic silencing in neuroscience applications. More broadly, PPO grants spatial and temporal control of G protein signaling cascades which may have additional use in dissecting fundamental aspects of cellular biology and GPCR pharmacology.

## RESULTS

### Spectral characterization of a photoswitchable GPCR-based opsin

Lamprey parapinopsin (PPO) is a non-visual GPCR-based opsin that interconverts between on and off states by absorption of ultra-violet (UV) and amber light, respectively (Figure 1A) (Koyanagi et al., 2004). Recently PPO was shown to couple to mammalian G_i/o_ proteins (Eickelbeck et al., 2019; Kawano-Yamashita et al., 2015), however the need for UV irradiation is a major limiting factor for use *in vivo*. We recently used violet light to stimulate PPO and control immune cell migration *in vitro* (O’Neill et al., 2018) and hypothesized that PPO may also be sensitive to even longer wavelengths, and thus suitable for *in vivo* applications.

**Figure 1.**
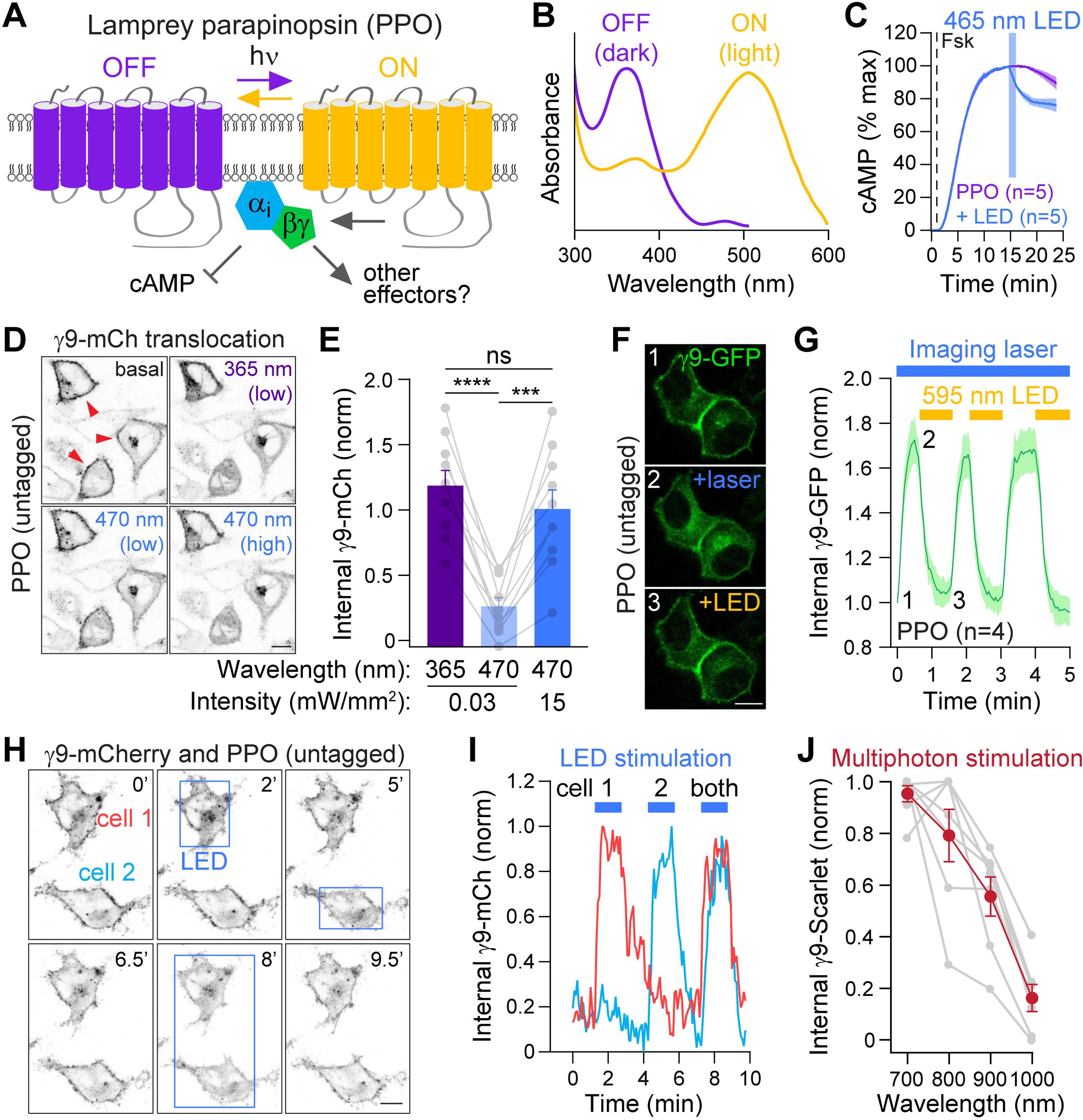
Spectral characterization of a photoswitchable GPCR-based opsin. (A) Cartoon of lamprey parapinopsin (PPO), a photoswitchable GPCR that is activated by UV light and turned off by amber light. PPO was previously shown to couple to Gi signaling pathways *in vitro*. (B) Absorption spectra of purified PPO protein in the dark off state (purple) and after irradiation with UV light (amber). Note the small hump in PPO absorbance in the off state in the blue light spectrum around 470 nm. Graph is modified from (Koyanagi et al., 2004) (copyright 2004 National Academy of Sciences, USA) (C) Optical stimulation with blue LED light (465 nm, 15.6 mW/cm^2^) inhibits forskolin-induced cAMP luminescence in PPO-expressing HEK cells (n= 5). (D) Live-cell images of Gβγ translocation assays using γ9-mCherry fluorescence to assess GPCR activation. In basal conditions, γ9-mCherry is localized to the plasma membrane (red arrows), but translocates after stimulation of PPO (untagged) with UV (365 nm) or blue (470 nm) light. Scale bar is 10 μm. (E) Quantification of γ9-mCherry translocation following stimulation with UV *vs.* blue light. Note that higher intensity blue light produces equivalent mobilization of Gβγ subunits to UV light. n=10 cells, Paired t-test, ns=not significant, ***p<0.001, ****p<0.0001. (F) γ9-GFP fluorescence in translocation assays to assess PPO photoswitching with amber light. γ9-GFP was imaged through a spinning disk every 3 seconds with a 488 nm laser (∼45 μW), which activated PPO and caused translocation from the plasma membrane (+laser). Simultaneous widefield illumination with a 595 nm amber LED (∼150 μW) resulted in γ9 translocation back to the membrane (+LED). Scale bar is 10 μm. (G) Quantification of γ9-GFP cytosolic translocation by the blue imaging laser and reversal with amber LEDs. Note the lack of desensitization and ability to repeatedly photoswitch PPO on and off. n=4 cells. (H) Individual photoactivation of cells expressing γ9-mCherry and untagged PPO. Blue boxes depict regions illuminated with blue LED light at different time-points, causing γ9-mCherry translocation from the membrane. Scale bar is 10 μm. (I) Individual traces of γ9-mCherry translocation from cells marked in panel H. Blue light triggered PPO activation at cellular resolution without inducing translocation in neighboring cells. (J) Quantification of the multiphoton spectra of PPO activation using γ9-translocation assays. γ9-mScarlet fluorescence was imaged with a Ti:Sapphire laser tuned to 1080 nm, while delivering multiphoton stimulation to PPO with a second laser at 700-1000 nm. n=7 cells.

We noticed a small peak in the blue wavelength range (450 – 500 nm) of the PPO absorption spectra (Figure 1B) which could greatly extend the utility of this opsin for neuroscience applications. We therefore tested if blue light sources commonly used in optogenetic experiments can activate PPO. We confirmed that purified PPO can indeed absorb non-UV light (**Figure S1A**). and found that blue LED stimulation (465 nm, 5 mW, 60s) inhibited forskolin-induced cAMP levels in HEK293 cells expressing PPO (Figures 1C and S1B). To more precisely define the spectral and temporal properties of PPO signaling, we used imaging assays of G protein translocation to monitor PPO engagement of downstream Gβγ subunits (Karunarathne et al., 2012; O’Neill et al., 2012). We co-expressed PPO with fluorescently tagged γ9 subunits in HeLa cells and quantified γ9 intracellular fluorescence changes from the plasma membrane. Imaging of mCherry-tagged γ9 with a 595 nm laser did not affect membrane localization (red arrows, Figure 1D), consistent with the action spectra of PPO that would keep the receptor in the off state (Figure 1B). Widefield illumination with low intensity UV light (365 nm, 0.03 mW/mm^2^, 150 ms pulse delivered every 5 s) produced rapid intracellular translocation of γ9-mCherry from the plasma membrane (Figure 1D **and** 1E; **Video S1**). Photoactivation with blue light of the same intensity (470 nm, 0.03 mW/mm^2^, 150 ms pulse every 5s) caused minimal translocation; however, when we increased the blue light intensity to 15 mW/mm^2^ we observed equivalent levels of translocation compared to UV illumination (Figure 1D **and** 1E). This suggests that while PPO exhibits lower blue light sensitivity, higher intensity stimulation produces equivalent efficacy. To examine the photoswitching properties of PPO, we imaged GFP-tagged γ9 subunits with a 488 nm blue laser (45 μW average power through the spinning disk, typical for GFP imaging), which activated PPO and increased the intracellular localization of γ9-GFP upon imaging (Figure 1F **and** 1G). Simultaneous widefield illumination with an amber LED (595 nm, 150 μW), rapidly reversed translocation of γ9-GFP back to the plasma membrane even while photostimulating PPO. We could repeatedly photoswitch PPO on and off without any apparent desensitization or changes in efficacy (Figure 1G). Thus, these data demonstrate that UV-sensitive PPO can also be activated by blue light and rapidly turned off with amber light, making it more suitable for use *in vivo*.

### Multiplexed optical GPCR activation and imaging

The lack of spontaneous γ9 translocation while imaging, and the ability to photoswitch PPO into the off state, suggested that PPO could be simultaneously employed with red fluorescent activity sensors. We first assessed PPO trafficking in cultured striatal neurons where we observed prominent localization at the membrane and distal dendrites (**Figure S1C**). To determine if we could utilize PPO for photoinhibition while monitoring neuronal activity, we co-expressed PPO with the genetically encoded Ca^2+^ indicator RCaMP1.07 (Ohkura et al., 2012) in striatal neurons. We observed spontaneous Ca^2+^ transients that were unaffected by imaging with a 561 nm laser. However, blue light stimulation of PPO-expressing neurons rapidly suppressed these transients and led to increased activity in adjacent PPO-negative neurons, possibly through disinhibition (**Movie S2).** To determine whether PPO could be used for more restricted patterns of G protein activation we imaged γ9-mCherry and then illuminated individual cells with blue light directed by a digital mirror device. We observed rapid translocation of γ9-mCherry in stimulated regions without altering membrane localization in adjacent cells (Figure 1H **and** 1I; **Video S3).** We were also able to achieve subcellular resolution of G protein activation (**Figure S1D and S1E**), indicating that PPO could represent a useful tool for probing the subcellular dynamics of GPCR signaling (Eichel and von Zastrow, 2018; Lobingier and von Zastrow, 2019).

We next asked whether this GPCR-based opsin could be activated by multiphoton stimulation. We imaged γ9-mScarlet translocation at 1080 nm, while photostimulating at different infrared wavelengths with a second tunable laser. Consistent with the single photon absorption spectra, we observed maximal G protein activation around 700 nm, which decreased at longer wavelengths tested up to 1000 nm (Figures 1J and **S1F**). While we did observe slight activation with the imaging laser at 1080 nm, we could reliably reverse these effects with amber LED light, similar to our single-photon activation experiments. Taken together, these molecular features suggest that PPO represents a unique optogenetic tool that could be used for simultaneous optical monitoring and activation of GPCR signaling cascades *in vivo* (Carrillo-Reid et al., 2019; Marshel et al., 2019; Packer et al., 2014; Rickgauer et al., 2014; Yang et al., 2018).

### PPO rapidly and reversibly inhibits neuronal voltage-activated calcium channels

To determine whether PPO has applications as an optogenetic silencing tool, we next generated a Cre-dependent (double-floxed inverted open reading frame, DIO) adeno-associated viral (AAV) construct for cell-specific expression (**Figure S2A**). We first examined PPO expression and trafficking in cultured dorsal root ganglia (DRG) neurons from *Avil*^Cre^ mice (da Silva et al., 2011; Zhou et al., 2010) transduced with AAVs to express Venus-tagged PPO (AAV5:CAG:DIO:PPO-Venus). Following transduction, we observed expression within 24 hours and found that PPO-Venus was trafficked along tau-labeled axons where it colocalized with the presynaptic marker synapsin-1 at terminals (Figure 2A).

**Figure 2.**
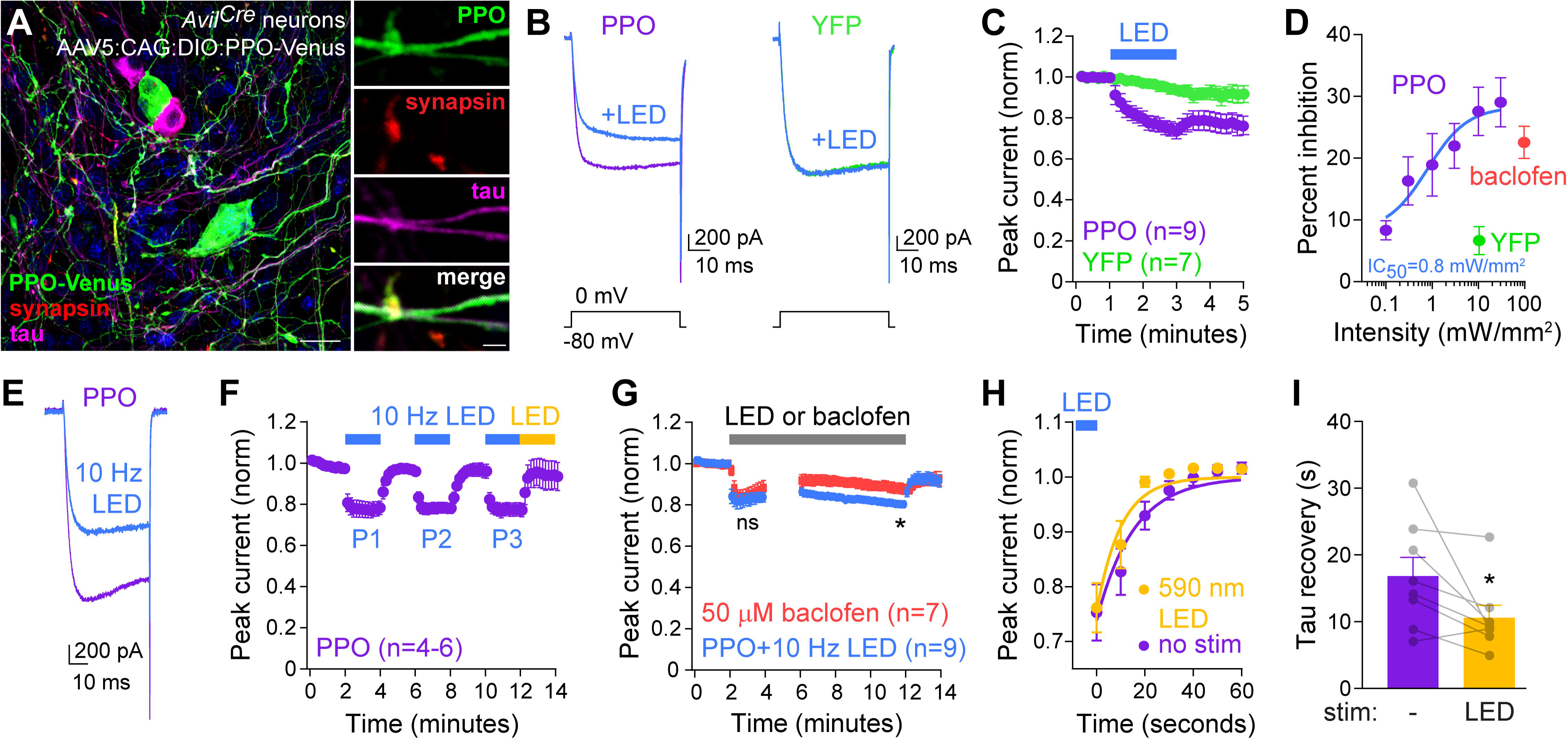
PPO inhibits neuronal calcium channel function. (A) Confocal micrograph of cultured DRG neurons from *Avil^Cre^* mice 7 days after transduction with Cre-dependent AAVs to express PPO-Venus (AAV5:CAG:DIO:PPO-Venus). The merged image shows PPO-Venus fluorescence (green) at cell bodies and tau^+^ axons (magenta). PPO is also expressed in synapsin-1^+^ axonal boutons (red). Scale bars are 20 μm (left) and 5 μm (right). (B) Ca^2+^ channel current traces elicited by voltage steps from −80 to 0 mV in cultured DRG neurons. Constant LED illumination (470 nm, 10 mW/mm^2^) inhibited currents in PPO-expressing neurons (purple) but not YFP^+^ controls (green). (C) Quantification of normalized Ca^2+^ channel currents depicting the time-course in inhibition in PPO^+^ (purple) *vs.* YFP control (green) neurons. Blue bar indicates when neurons were stimulated with constant blue LED light (470 nm, 10 mW/mm^2^). (D) Dose-response curve of Ca^2+^ channel inhibition in response to varying intensities of constant blue light to PPO+ neurons (purple) or YFP+ controls (green). The inhibitory-half maximal value (IC_50_) of 0.8 mW/mm^2^ was calculated by a non-linear fit of the responses. Inhibition by the GABA_B_R agonist baclofen (50 μM, coral) was used to compare efficacy of PPO stimulation. n=6-11 for PPO at each intensity. n=7 for YFP controls and n=20 for baclofen responses. (E) Ca^2+^ channel trace of PPO expressing neuron before (purple) and after (blue) stimulation with pulsed blue light (470 nm, 10 Hz, 10 ms pulse-widths, 10 mW/mm^2^). (F) Time course of peak current inhibition to repeated LED pulses (blue bars) and enhanced recovery in amber light (yellow, 590 nm LED). n=4-6 neurons. (G) Plot of normalized Ca^2+^ channel currents in response to prolonged stimulation of PPO with 10 Hz LED light (blue) or GABA_B_R activation with 50 μM baclofen (coral). PPO and GABA_B_Rs exhibited similar levels of short-term inhibition (ns=not significant); however, the response to baclofen desensitized within several minutes, while PPO continued to inhibit Ca^2+^ channels without apparent desensitization. T-test, *p<0.05 (H) Normalized recovery time-course of Ca^2+^ channel currents with (LED, amber) and without (no stim., purple) subsequent illumination by amber light. The blue bar depicts when the pulsed LED was turned off. n=8 cells for each. (I) Summary graph of the plotted recovery tau with (amber) and without (purple) subsequent illumination with a 590 nm LED to photoswitch PPO off. Paired t-test, *p<0.05

To examine PPO coupling to neuronal G protein signaling cascades, we next recorded voltage gated Ca^2+^ channel currents in DRG neurons, where pharmacological activation of endogenous G_i_-coupled GPCRs inhibits Ca^2+^ currents through Gβγ-mediated voltage-dependent mechanism (Bean, 1989; Bourinet et al., 1996; Currie, 2010; Dunlap and Fischbach, 1981). Constant illumination of PPO-expressing dorsal root ganglion (DRG) neurons with 470 nm LED light rapidly suppressed calcium channel currents in a light intensity-dependent manner (Figure 2B-D). Maximal inhibition plateaued around 10 mW/mm^2^ and was equivalent to that observed with the GABA_B_ receptor agonist baclofen (Figure 2D). Blue LED light inhibited DRG calcium channel currents with an IC_50_ of 0.8 mW/mm^2^, and no inhibition was observed in YFP-expressing controls (10 mW/mm^2^, Figure 2B-D).

Surprisingly, in contrast to our Gβγ translocation assays, Ca^2+^ channel currents did not spontaneously recover after constant blue light illumination at higher intensities (Figures 2C and S2B), and we were unable to reverse this effect by photoswitching PPO with green or amber LEDs (**Figure S2C**). Shorter durations of constant photostimulation (10 mW/mm^2^) did however recover (data not shown). This dose- and time-dependent irreversible inhibition suggested that we might be photobleaching the opsin with constant illumination at higher light intensities, so we tested whether agonist pulses of blue light could be used to activate PPO in a readily reversible manner. We explored a variety of stimulation parameters and found that 10 ms pulses delivered at 10 Hz with 10 mW/mm^2^ intensity exhibited similar efficacy to the maximal inhibition observed with constant illumination, but currents recovered rapidly to baseline (Figure 2E **and** 2F). We also tested the temperature dependence of Ca^2+^ channel inhibition and reversal, which were similar at physiological and room temperature (**Figure S2D and S2E**). We could repeatedly inhibit currents with similar efficacy (Figures 2F and S2F) suggesting that PPO undergoes little functional desensitization. To more rigorously test this, we compared Ca^2+^ channel inhibition from prolonged optical stimulation to activation of endogenous GABA_B_Rs with baclofen, a well-established G_i_-coupled inhibitor of neurotransmitter release at synaptic terminals. We observed similar degrees of inhibition between PPO and GABA_B_R after 2 minutes, however the baclofen response desensitized ∼50% over the course of 10 minutes, while PPO did not exhibit any observable functional desensitization (Figure 2G **and S2G**). Consistent with our reversal of Gβγ localization, we were able to speed the recovery of Ca^2+^ channel inhibition approximately two-fold by illuminating neurons with 590 nm LED light (Figure 2H **and** 2I). The recovery tau of ∼10 seconds reflects both the photoconversion of PPO to the off state and G protein deactivation, with the latter likely to be the limiting step in terminating PPO signaling.

While we did not observe functional desensitization, illumination of PPO with UV light induces translocation of lamprey β-arrestin, which was proposed to internalize receptors and terminate signaling similar to mammalian rhodopsins (Kawano-Yamashita et al., 2011; Wilden et al., 1986). To determine whether PPO undergoes arrestin-dependent internalization and desensitization following blue light activation in mammalian cells, we performed live cell total internal reflection fluorescence (TIRF) microscopy of GFP-tagged arrestin-3 and mCherry-tagged clathrin to mark endocytic zones. Illumination of PPO-expressing cells induced arrestin clustering at these clathrin^+^ sites (**Figure S2H and S2I**), confirming that PPO activation can activate mammalian arrestins. However, PPO did not accumulate with arrestins at these sites (**Figure S2J**), and we did not observe any internalization of PPO even after 90 minutes of stimulation with pulsed blue light (**Figure S2K**). Thus, while PPO activation clearly engages and mobilizes arrestin, we did not detect receptor internalization or functional desensitization in both imaging and physiological measures, which has been observed with other types of GPCRs (Eichel and von Zastrow, 2018; Eichel et al., 2016, 2018; Nuber et al., 2016). Collectively, these data demonstrate that pulsed blue light stimulation of PPO can be used to rapidly and reversibly inhibit neuronal calcium channel function, with equivalent efficacy to endogenous G_i_-coupled GABA_B_Rs.

### Calcium channel inhibition is G protein dependent

We next asked whether PPO inhibition of Ca^2+^ channel currents was G protein mediated. Treatment of DRG neurons with pertussis toxin (PTx) to inactivate Gα_i/o_ subunits and inhibit G protein signaling (Dolphin and Scott, 1987; Katada and Ui, 1982; Scott and Dolphin, 1987) completely blocked Ca^2+^ channel inhibition by both blue LED stimulation of PPO and activation of endogenous GABA_B_Rs with baclofen (Figure 3A-C). While this mechanism is consistent with PPO activation of Gα_i_ signaling, we also inquired whether PPO couples to Gα_o_ subunits. To test this, we performed γ9 translocation assays in cell treated with PTx to inactivate Gα_i/o_ proteins and and co-expressed a PTx-resistant Gα_o_ mutant (Hunt et al., 1994; Wise et al., 1997). PTx treatment prevented translocation of γ9-mScarlet during blue light stimulation of PPO-expressing cells; however, expression of PTx-resistant Gα_o_ subunits partially restored γ9 translocation (**Figure S3A and S3B**), demonstrating that PPO can also couple to Gα_o_ signaling pathways.

**Figure 3.**
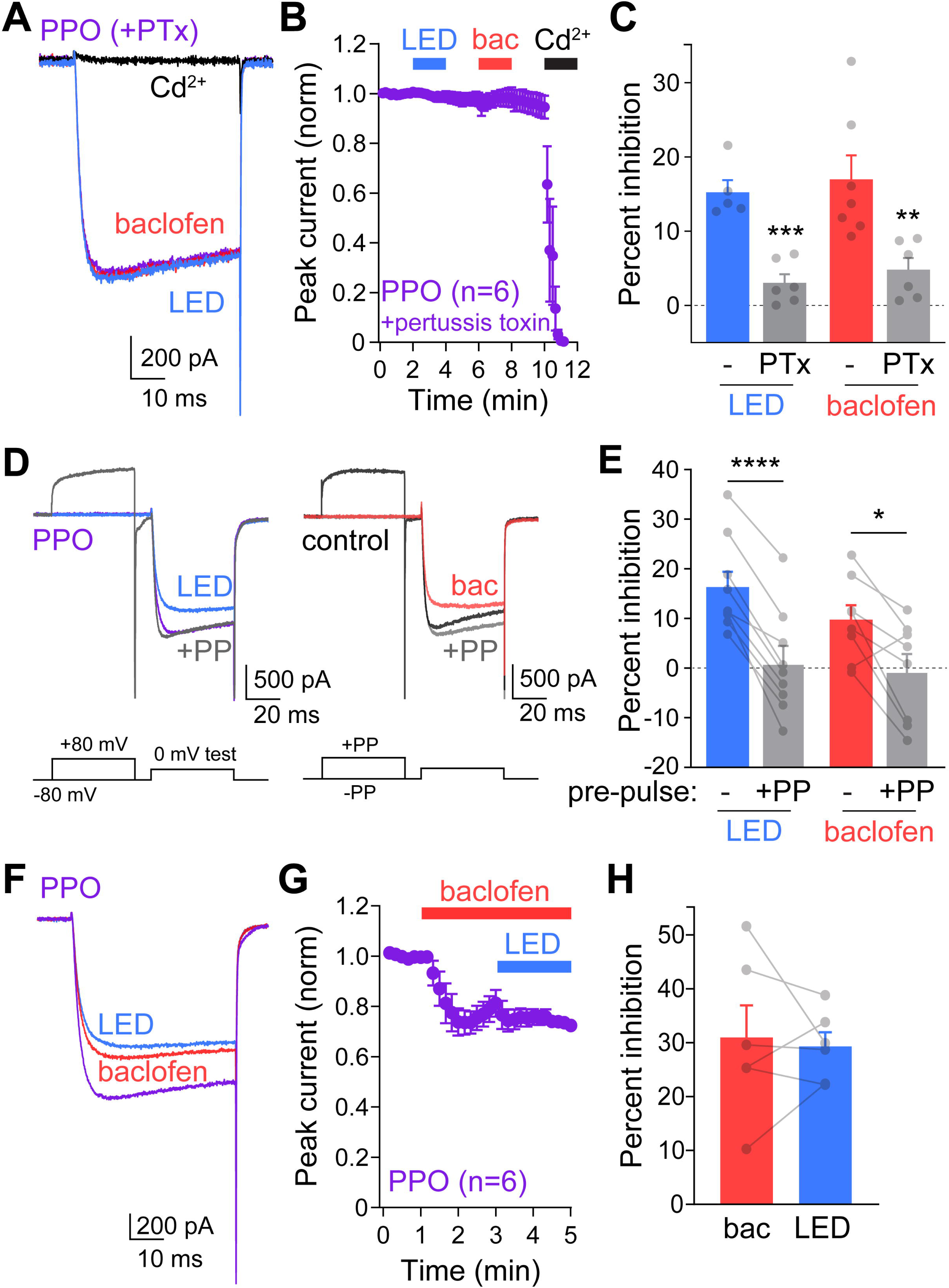
PPO couples to neuronal G proteins. (A) Ca^2+^ channel trace of PPO expressing neuron treated for 24 hours with 200 ng/ml pertussis toxin (PTx, purple) and subsequent responses to 10 Hz blue light (blue), 50 μm baclofen (coral) and 100 μm Cd^2+^ (black) to block Ca^2+^ channels. (B) Time course of normalized peak Ca^2+^ channel currents in PPO-expressing neurons treated with PTx (purple) in response to 10 Hz LED stimulation (blue), 50 μm baclofen (coral), and 100 μm Cd^2+^ (black). (C) Summary graph of the percent inhibition of Ca^2+^ channels in PPO expressing neurons. The blue and red bars represent parallel controls that were not treated with PTx. Gray bars represent PTx-treated PPO^+^ neurons stimulated with blue light (left) or baclofen (right). n=5-7 for each, t-test **p<0.01, ***p<0.001. (D) Ca^2+^ channel current traces +/− pre-pulse (PP) depolarizations to 80 mV before test pulses to 0 mV to test for voltage-dependent inhibition. Traces depict PPO (purple) or control (black) cells without pre-pulse, then after LED illumination of PPO (blue) or baclofen (coral). 50 ms pre-pulse depolarizations (gray) relieved the inhibition by LED (left) or baclofen (right). (E) Summary graph of voltage-dependent inhibition by LED illumination of PPO-expressing cells (n=9), or control neurons stimulated with baclofen (n=8). In both cases, inhibition was completely relieved by the pre-pulse. Paired t-test ****p<0.0001, *p<0.05. (F) Ca^2+^ channel trace of PPO expressing neuron (purple) stimulated with 50 μM baclofen (coral) followed by 10 Hz LED stimulation (blue). (G) Time course of normalized peak Ca^2+^ channel currents in PPO-expressing neurons (purple) stimulated by 50 μM baclofen (coral) and 10 Hz LED pulses (blue). n=6 (H) Summary graph of the percent inhibition of Ca^2+^ channel currents by baclofen and subsequent LED stimulation. n=6

Following activation of G_i_-coupled GPCRs, the Gβγ subunit is the primary downstream modulator of channel function in neurons (Lüscher and Slesinger, 2010; Zamponi and Currie, 2013). We tested whether PPO mobilized Gβγ subunits to directly inhibit Ca^2+^ channels through voltage-dependent mechanism (Bean, 1989; Herlitze et al., 1996; Ikeda, 1996). Strong depolarizing pre-pulse steps reversed Ca^2+^ channel inhibition by PPO and baclofen (Figure 3D **and** 3E), while subsequent I-V curves revealed no change in gating (**Figure S3C-E**). Thus, light stimulation of PPO is consistent with the well-established voltage-dependent mechanism of Ca^2+^ channel inhibition by G_i_-coupled GPCRs. Our previous development of opto-XRs has suggested that rhodopsin-based chimeras couple to endogenous and overlapping GPCR signaling cascades (Siuda et al., 2015). To test whether PPO converges with endogenous G_i_-coupled GPCRs, we recorded Ca^2+^ channel currents and activated GABA_B_Rs with baclofen. Ca^2+^ channel inhibition plateaued within ∼1 minute and then desensitized slightly (Figure 3F **and** 3G). Subsequent illumination of PPO with blue light pulses did not increase the peak inhibition by baclofen (Figure 3F-H). Thus, PPO couples to the same G protein signaling cascades in neurons as native G_i_-coupled GABA_B_Rs to inhibit voltage-gated calcium channels.

### Reversible silencing of synaptic transmission

Presynaptic G_i/o_-coupled GPCRs, like GABA_B_Rs, exert powerful control over synaptic transmission by inhibiting vesicle release through multiple mechanisms, including decreased calcium channel function and interference with release machinery (Blackmer, 2001; Browning et al., 2002; Heinke et al., 2011; Stachniak et al., 2014; Takahashi et al., 1998; Zurawski et al., 2019a). The indistinguishable G_i/o_ coupling between PPO and endogenous GABA_B_Rs, suggested that PPO might represent a unique optical approach for presynaptic inhibition that takes advantage of these endogenous silencing mechanisms. We next tested whether PPO alters synaptic transmission in excitatory thalamic projections to the barrel cortex, which was used previously to identify biophysical limitations of optical inhibition at terminals with Cl^-^ and H^+^ pumps (Mahn et al., 2016). We injected Cre-dependent AAVs carrying PPO-Venus into the ventral posterior medial nucleus (VPM) of the thalamus in *Vglut2-IRES-Cre* mice to target glutamatergic neurons (Figure 4A) (Vong et al., 2011). We first assessed PPO trafficking along the axons and into terminals of this long-range projection using confocal microscopy with deconvolution. We observed PPO-Venus in thalamocortical axons innervating the barrel cortex in as little as 14 days after injection, similar to the rapid trafficking of ChR2 observed previously (Crandall et al., 2017), with dense projections to layer IV neurons (Figure 4B). Imaging of these terminals revealed colocalization of PPO-Venus with the presynaptic glutamate transporter vGluT2 which was opposed to excitatory synapses marked by PSD-95 (Figure 4C). These experiments demonstrate that PPO traffics effectively into axon terminals *in vivo*, making it well-positioned to control presynaptic release.

**Figure 4.**
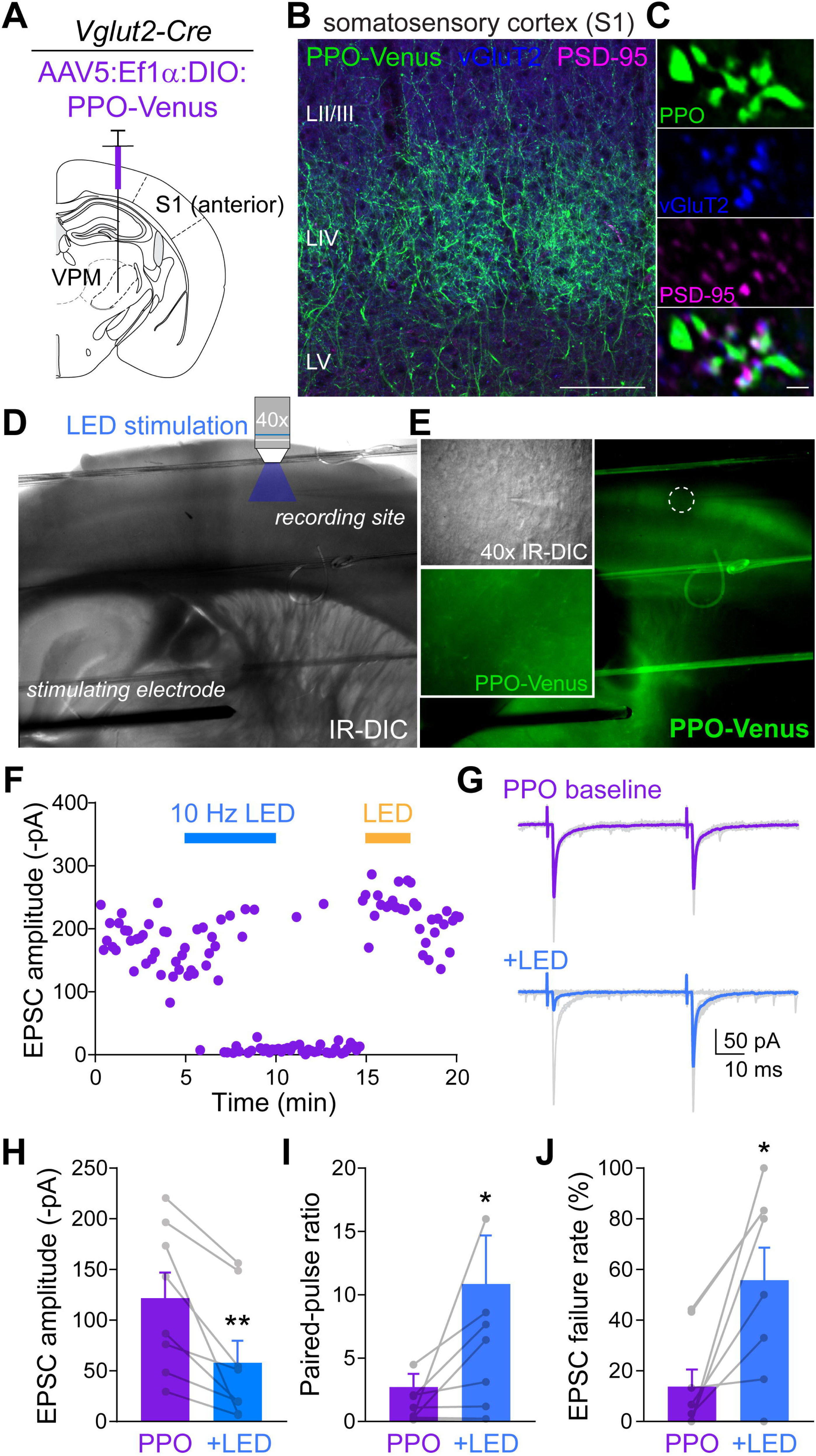
Presynaptic PPO can silence synaptic transmission. (A) Cartoon depicting the injection site of Cre-dependent AAVs (AAV5:Ef1α:DIO:PPO-Venus) into the ventral posterior medial nucleus (VPM) of the thalamus in *Vglut2-Cre* mice to target excitatory projections to primary somatosensory cortex (S1). (B) Confocal micrograph of the somatosensory cortex from a *Vglut2-Cre* mouse injected with Cre-dependent AAV5s expressing PPO-Venus in the VPM. PPO-Venus^+^ axons (green) can be seen predominantly in layer IV of the barrel cortex. vGluT2+ presynaptic terminals (blue) and the excitatory postsynaptic marker PSD-95 (magenta) were used to visualize the synaptic localization of PPO. Scalebar represents 100 μm. (C) Deconvolved super-resolution image depicting colocalization of PPO (green) with presynaptic vGluT2+ terminals (blue) opposed to excitatory postsynaptic sites marked by PSD-95 (magenta). Scale bar represents 1 μm. (D) IR-DIC images of an acute thalamocortical slice depicting the location of the bipolar stimulating electrode in the VPM, and the patch-pipette and objective for optical stimulation in LIV of the barrel cortex. (E) Epifluorescence image of PPO-Venus is the same acute slice showing PPO trafficking along axons projecting to the barrel cortex. Insets show 40x IR-DIC images of a recorded neuron in LIV surrounded by PPO-Venus^+^ fibers. PPO-Venus fluorescence was intentionally bleached (dashed circle) to show that the area of photostimulation was confined to individual barreloids in layer IV. (F) Plot of electrically evoked EPSCs which were silenced by 10 Hz pulsed LED stimulation (blue bar, 470 nm) and reversed by subsequent illumination with amber light (590 nm). (G) Representative EPSC traces (gray) and averaged responses during baseline (purple) and 10 Hz LED stimulation (blue). Paired pulses were delivered at 50 ms intervals. Note the increased number of synaptic failures after LED stimulation. (H-J) Summary graph of EPSC amplitudes (H) paired pulse ratios (I), and failure rates (J) before (purple) and after LED stimulation (blue). n=8, paired t-tests, *p<0.05, **<0.01.

We assessed PPO at this long-range thalamocortical slice preparation where postsynaptic neurons in cortex receive inputs from intact axonal collaterals originating several millimeters away in thalamus (Agmon and Connors, 1991; Cruikshank et al., 2007). This permits electrical stimulation of cell bodies in the VPM while constraining the photostimulation to terminals in layer IV of the barrel cortex, limiting a potential confound of light “spillover” and somatic activation in shorter projections (Figure 4D). In acute slices prepared from mice receiving viral injections only 2 weeks prior, we observed PPO-Venus expression along the entire axonal projection and at terminals surrounding layer IV cortical neurons (Figure 4E). We electrically evoked monosynaptic excitatory synaptic currents in VPM, and then photostimulated PPO-expressing terminals through the microscope objective (Figure 4D-G). Blue light pulses (10 Hz, 10 ms, 10 mW/mm^2^) inhibited EPSC amplitudes, and in some neurons completely silenced synaptic transmission, which could be reversed by constant illumination with amber light (Figure 4F-H). Consistent with a presynaptic mechanism of inhibition previously reported for numerous G_i/o_-coupled GPCRs (Johnson and Lovinger, 2016; Kreitzer and Malenka, 2007; Nanou and Catterall, 2018; Zucker and Regehr, 2002; Zurawski et al., 2019b), paired-pulse ratios and failure rates were increased (Figure 4I **and** 4J). These data collectively demonstrate that optical stimulation of PPO reversibly couples to inhibitory GPCR signaling cascades to inhibit presynaptic vesicle release at long-range excitatory projections in the brain.

### PPO is functional in vivo, and inhibits dopamine-dependent reward-seeking behaviors

To functionally determine whether PPO can be employed as an inhibitory opsin for *in vivo* use, we first assessed if PPO could suppress reward-seeking behaviors mediated by dopamine (DA) neurons in the ventral tegmental area (VTA) (Morales and Margolis, 2017; Parker et al., 2019). We bilaterally injected Cre-dependent AAVs into *DAT-IRES-Cre* mice (Bäckman et al., 2006) to express PPO in DA neurons of the VTA, and implanted optical fibers above their somata (Figures 5A). We confirmed that PPO was strongly expressed throughout the VTA and found that it trafficked very well in DA neurons along cell bodies and processes (Figures 5B and **S4A and S4B**). Reward-seeking behavior was tested in mice trained to perform operant tasks for a sucrose reward (Figure 5C-E). Food-restricted mice underwent Pavlovian conditioning to associate a cue (house light) with access to a sucrose sipper (Figure 5D) and were trained on fixed ratio (FR) schedules to nose poke for sucrose rewards in response to the cue (Figure 5E). We then assessed reward seeking behavior in FR-3 tests with and without optical inhibition. 10 Hz laser light (473 nm, 10 ms pulses) at the cell bodies of VTA DA neurons significantly suppressed both the number of nose pokes and rewards (Figure 5F). These data indicate reduced reward-seeking behavior when VTA DA neurons were inhibited by PPO in the soma and dendrites of these neurons, consistent with other inhibitory approaches (Corre et al., 2018; Tye et al., 2013). In addition, these data also suggest that this GPCR-based inhibitory approach may have potential advantages over many ion-conducting or pumping opsins, given that PPO can inhibit somatic activity in a pulsed mode without potential confounds of constant illumination (Stujenske et al., 2015).

**Figure 5.**
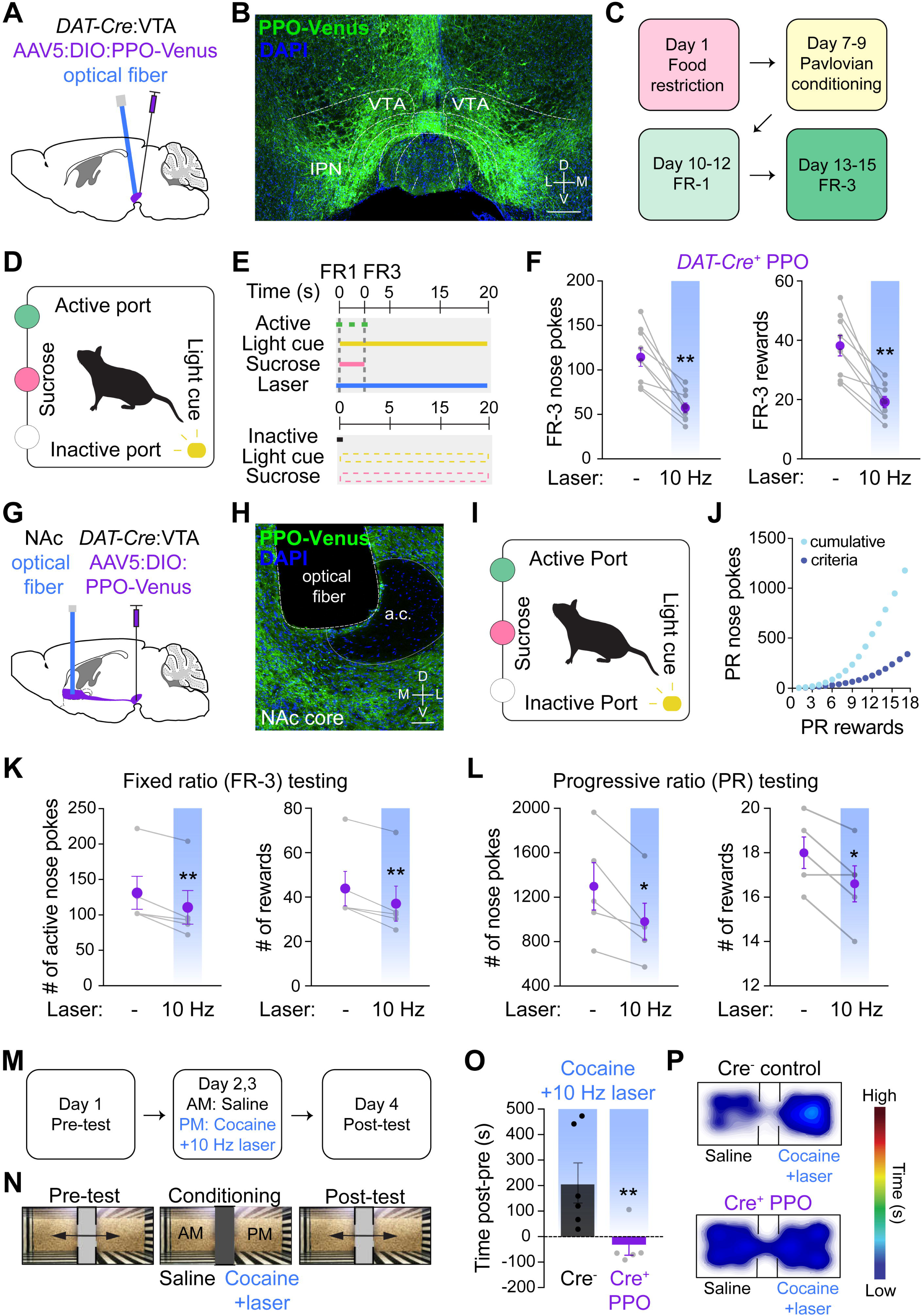
PPO inhibits dopamine neuron terminals to suppress reward seeking behaviors. (A) Cartoon depicting the experimental strategy for expressing Cre-dependent PPO-Venus (AAV5:Ef1α:DIO:PPO-Venus) in *DAT-Cre+* dopamine neurons in the ventral tegmental area (VTA). Optical fibers were implanted bilaterally above the VTA to stimulate cell bodies. (B) Confocal micrograph demonstrating PPO-Venus expression in cell bodies of the VTA. Scale bar = 400 μm. (C) Experimental timeline for operant task training for sucrose rewards to assess reward seeking behaviors. Mice were trained on fixed ratio (FR)-1 and FR-3 schedules. (D) Cartoon of an operant chamber in which mice nose poke into an active port in response to a light cue to receive a sucrose reward. (E) Experimental design of the operant task in which mice learn to nose poke in the active port for a sucrose reward in response to the light cue. No reward is given for nose pokes in the inactive port. During FR1 training mice must nose poke once, while FR3 training requires 3 nose pokes, for a sucrose reward. (F) Summary graphs operant behaviors with and without laser stimulation of PPO-expressing VTA neurons. Blue columns indicate pulsed laser light (473 nm, 10 Hz, 10 ms, 5-8 mW) which decreased both the number of nose pokes (left) and rewards (right). n=9 mice each, paired t-test **p<0.01 (G) Cartoon depicting sites for inhibiting DA neuron projections to the nucleus accumbens (NAc). Cre-dependent AAVs were injected into the VTA of *DAT-Cre* mice and optical fibers were placed above their terminals in the NAc shell. (H) Confocal micrograph depicting PPO-Venus expression in DA neuron terminals of the NAc. The placement of the optical fiber for targeting these projections is shown. Scale bar = 100 μm. (I) Operant chamber cartoon, as described above. (J) Example graph of showing the escalating number of nose pokes needed to receive a reward in progressive ratio (PR) testing. (K) Summary graphs of FR-3 testing with and without optical stimulation of DA terminals as above (blue bars). Terminal inhibition decreased the number of rewards that mice received (right) but not the number of nose pokes (left). n=5 mice. Paired t-test **p<0.01 (L) Summary graphs of PR testing in which optical stimulation decreased both the number of nose pokes and rewards received, indicating reduced motivation to work for these rewards when DA terminals to the NAc were inhibited. Paired t-test *p<0.05 (M,N) Experimental timeline and set-up for testing cocaine preference behaviors in mice. During pre-testing mice were allowed to freely explore the 2 chambers. During pairing, mice were given saline injections in 1 chamber and cocaine (amount) along with 10 Hz blue light to inhibit DA neuron terminals in the other chamber. After 2 days of pairing, preference for either chamber was determined in both Cre+ PPO and Cre-control mice. (O) Summary graph of difference in time spent in the cocaine-paired chamber after testing. Cre-control mice (n=6) exhibit strong preference for the cocaine-paired chamber which was completely blocked by blue laser stimulation of DA terminals in the NAc in PPO-expressing mice (n=5). Both groups received optical stimulation during cocaine-pairing. t-test **p<0.01 (P) Heat maps of relative time spent in each chamber for Cre-negative control mice (top) and PPO-expressing mice (bottom).

To determine the efficacy of PPO in silencing DA neuron *projections* to the NAc, we used the same strategy to virally target VTA DA neurons, and implanted bilateral optical fibers above their terminals in the NAc core (Figure 5G). We observed strong PPO expression in VTA neuron terminals in the NAc (Figures 5H and S4B), consistent with the efficient trafficking of PPO we observed in all other diverse neuron types examined (glutamate, GABA, DA and DRG). We performed similar operant tasks for sucrose rewards in these mice, but also included progressive ratio (PR) tests (Figure 5I-J). In these PR tests, the number of nose pokes required to receive a reward was escalated exponentially, which assesses the point at which the animal is no longer willing to work for a reward (Figure 5J) (Hodos, 1961; Parker et al., 2019; Richardson and Roberts, 1996). Mice underwent similar training schedules before FR and PR testing. Photoinhibition of DA projections to the NAc core suppressed active total nose poke behavior in FR testing and reduced the number of rewards obtained (Figure 5K). In PR testing, optical stimulation depressed both nose pokes and the number of rewards in Cre^+^ mice expressing PPO (Figures 5L and S4C) but not in Cre^-^ controls (**Figure S4D**), consistent with the role of this DA projection in both reward-seeking and motivated behaviors (Corre et al., 2018; Tye et al., 2013).

We finally tested whether PPO could inhibit behavioral states of increased dopaminergic tone by examining cocaine-induced conditioned place preference. We used the same approach as above to optically target these terminals in *DAT-*Cre^+^ mice and Cre negative controls (Figure 5G). After pre-testing, mice were paired with saline injections in one chamber followed by cocaine pairing in the other chamber (Figure 5M **and** 5N). During cocaine pairing however, both groups also received 10 Hz optical illumination of VTA DA terminals in the NAc. After 2 days of pairing, we assessed preference behavior for the two chambers. Cre^-^ control mice strongly preferred the cocaine-paired chamber, which was completely blocked in *DAT-*Cre^+^ mice expressing PPO (Figure 5O-P). Importantly, distances traveled were similar between the two groups (**Figure S4E**), demonstrating the selectively of PPO inhibition of DA terminals in the NAc and not collateral projections to dorsal striatum. Collectively, these data demonstrate that PPO is a useful and efficient optogenetic tool for photoswitchable control of inhibitory G protein signaling cascades to inhibit presynaptic terminals *in vivo*.

## DISCUSSION

Here we identify and characterize PPO as a novel photoswitchable G_i/o_-coupled GPCR for silencing presynaptic terminals. PPO rapidly couples to neuronal G proteins when illuminated with blue light and can be switched off with amber light. PPO is trafficked very efficiently in neurons to both distal dendrites and presynaptic terminals. Here, PPO can be used to silence glutamatergic transmission. Similarly, at dopamine neuron projections to the NAc, PPO suppressed reward-seeking behaviors and completely blocked cocaine place-preference. Thus, this photoswitchable GPCR fills a major gap in biology, pharmacology, and neuroscience fields, providing a unique optical tool for probing GPCR dynamics that can also achieve rapid and reversible synaptic silencing (Lin et al., 2013; Liu et al., 2019; Mahn et al., 2016; Wiegert et al., 2017). Indeed, the strength of this approach for synaptic silencing was independently demonstrated in a parallel study from the Yizhar lab using OPN3, a bistable G_i/o_-coupled opsin from mosquitos (Mahn et al., 2020). Furthermore, these opsins may represent developmental scaffolds for constructing new and improved photoswitchable opto-GPCR chimeras and for gaining spatiotemporal precision over traditional chemogenetic DREADD-based approaches.

### Spectral properties of PPO and technical considerations

PPO was first identified as the UV sensitive pigment in lamprey pineal cells (Koyanagi et al., 2004). Despite the optogenetic potential of a photoswitchable GPCR-based opsin, PPO seemed limited by the requirement for UV irradiation (Eickelbeck et al., 2019; Kawano-Yamashita et al., 2015). We took advantage of a peculiar shift in the activation spectrum and used blue laser and LED light sources, readily available in many biology labs, to stimulate PPO at light intensities comparable to those used in typical optogenetic experiments. This was important, as we were able to seamlessly integrate PPO into all of our optogenetic setups without needing to purchase any additional specialized equipment. Also, while many GPCR-based opsins possess exquisite light sensitivities that that can be problematic due to activation by ambient light, we were able to use pulsed light intensities commonly used for ion-conducting opsins like ChR2 (Mattis et al., 2011), by working at these “off-peak” wavelengths. Moreover, we observed that PPO is kept in the off state by ambient light (data not shown), limiting our initial concerns of unintended activation or constitutive activity prior to, or during, experiments. It is important to note that not all blue light sources are the same, particularly when working with LEDs. In some cases, their broad emission spectra may include UV or green photons, which could switch PPO into unintended states. For example, we could photoswitch PPO off with longer wavelength amber LEDs and even coherent 515 nm laser light but saw activation with 530 nm LEDs, owing to their broad emission (data not shown).

Stimulation with UV light appears to constitutively activate PPO until subsequent illumination with longer light wavelengths (Eickelbeck et al., 2019; Kawano-Yamashita et al., 2015). In contrast, the pulsed blue light that we used here seems to shift PPO into a more dynamic state in which G protein signaling deactivates ∼20 seconds after illumination. While this transition can be sped up by amber light (Figure 2G, 2H), the active state induced by pulsed blue light could reflect intermediate conformational states (Ernst et al., 2014; Kouyama and Murakami, 2010) which has recently been demonstrated with pharmacological activation of GPCRs (Du et al., 2019; Kato et al., 2019; Wingler et al., 2019). We find this possibility particularly intriguing, as it suggests that PPO coupling to G proteins could be dynamically altered with different durations or colors of “agonist” or light pulses, something not typically achieved with traditional pharmacological approaches. Thus, dynamic control of the G protein coupling state may allow for differential control of downstream effectors in future iterations of this GPCR-based optical tool. This could be used to alter signaling selectivity, akin to GIRK channel activation by the dynamics of M2 muscarinic, but not β2 adrenergic, receptor-coupling to release Gβγ subunits (Touhara and Mackinnon, 2018). This may also have important considerations for whether coupling to inhibitory signaling cascades has equivalent efficacies in different cell types.

### In vivo application of GPCR-based inhibitory opsins

PPO couples to mammalian Gα_i/o_ proteins to engage the canonical downstream cascades to inhibit adenylyl cyclase (Kawano-Yamashita et al., 2015), activate GIRK channels (Eickelbeck et al., 2019), and inhibit Ca^2+^ channels (this study). We focused on the latter mechanism to test whether PPO could fill a major gap in the systems neuroscience toolbox – that is, rapid and reversible inhibition of transmitter release from terminal projections (Wiegert et al., 2017). PPO rapidly and reversibly inhibited neuronal voltage-gated calcium channels with equal efficacy to the GABA_B_R agonist baclofen (Figure 2), which strongly suppresses synaptic transmission throughout the nervous system. While decreased Ca^2+^ influx by presynaptic G_i/o_-coupled GPCRs is a primary mechanism for inhibiting synaptic transmission, it is possible that PPO may also activate Gβγ subunits to directly inhibit SNARE proteins (Blackmer, 2001; Zurawski et al., 2019b). In the hippocampus, different types of presynaptic GPCRs can inhibit synaptic transmission through distinct mechanisms, with GABA_B_Rs reducing vesicle release primarily through Ca^2+^ channel inhibition while serotonin 5-HT_1B_ receptors exert SNARE-dependent effects (Zurawski et al., 2019a). When implementing PPO for *in vivo* behavior studies, it will be critical to assess how PPO affects synaptic transmission at the different projections being targeted, as presynaptic GPCRs can modulate synapses in an input specific manner, with varying efficacies (Burke et al., 2018; Creed et al., 2016; Kreitzer and Malenka, 2007; Marcott et al., 2018; Tejeda et al., 2017). Whether PPO can couple to distinct signaling cascades in different cell types or mimics presynaptic GPCRs to regulate plasticity certainly warrants future investigation.

The unique optical properties of PPO make it particularly intriguing for *in vivo* applications combining optogenetic manipulations with readouts of circuit activity (Carrillo-Reid et al., 2019; Marshel et al., 2019; Packer et al., 2014; Rickgauer et al., 2014; Yang et al., 2018). These approaches have typically used excitatory channel-based opsins together with genetically encoded Ca^2+^ indicators (GECIs) to test the contributions of specific cells within a circuit to a behavioral task (Carrillo-Reid et al., 2019; Marshel et al., 2019). By incorporating PPO into these experimental paradigms, it may now be possible to determine how inhibitory neuromodulator signaling influences circuit dynamics with more precise cellular and temporal resolution. PPO can be activated by multiphoton wavelengths up to ∼1100 nm, so red-shifted GECIs with excitation wavelengths exceeding 1200 nm (Dana et al., 2016; Inoue et al., 2019) could be used to limit PPO activation during *in vivo* imaging. This may open new opportunities for all-optical cell-specific manipulations and readouts of GPCR action across diverse areas cellular biology, pharmacology, and neuroscience fields.

### Conclusions and future directions

In its current state, PPO is an effective tool for rapid and reversible control of inhibitory GPCR signaling cascades. By coupling to endogenous inhibitory mechanisms, PPO is effective both at inhibiting cell bodies, likely through GIRK channel activation, and presynaptic inhibition of vesicle release. However, PPO may also represent a new chromophore scaffold for engineering photoswitchable chimeras to mimic the biological signaling of diverse GPCRs with spatiotemporal precision (Airan et al., 2009; Gunaydin et al., 2014; Li et al., 2015; Oh et al., 2010; Siuda et al., 2015; van Wyk et al., 2015). These molecular engineering efforts may require a more detailed understanding of the structural dynamics of PPO photoswitching, which in turn may also lead to variants with more rapid kinetics or tuning of the spectral properties. Another intriguing avenue is the generation of photoswitchable DREADD-like tools, for more precise spatial and temporal control over these powerful chemogenetic approaches for investigating neural circuits (Roth, 2016). More broadly, achieving precise optical control over different types of GPCRs may enable new studies into fundamental aspects of these clinically important drug targets, including functional selectivity, subcellular coupling, and signaling dynamics *in vivo*.

## Acknowledgements

This work was supported by a NIH BRAIN Initiative grant R01 MH111520 (MRB and RKS), NIH R21 DA049569 (BAC and PRO), NIH K01 DA042219 (PRO), NIH K01 DK115634 (VKS), and NIH R35 GM122577 (NG). We would like to thank Akihisa Terakita and Mitsumasa Koyanagi for initial advice in working with parapinopsin, Yu-Qing Cao for advice in recording calcium channel currents, and Ben Suter for suggestions in building photostimulation systems. We would also like to thank Fletcher Austin and Paul Gaetano for insightful discussions. This work was also supported by the Hope Center Viral Vectors Core at Washington University School of Medicine.

## Author contributions

Conceptualization: BAC, PRO, RWG, MRB

Methodology: BAC, PRO, RG, JJY, XM, KEP, VKS, RKS RWG, MRB

Investigation: BAC, PRO, RG, JJY, XM, KEP, SMS, AMV, AJE, VKS

Resources: MCS, VK, SKV

Writing – original draft: BAC

Writing – review and editing: BAC, PRO, RWG, MRB

Visualization: BAC, PRO, RG, RWG, MRB

Supervision: NG, RKS, RWG, MRB

Funding Acquisition: BAC, PRO, NG, RKS, RWG, MRB

## METHODS

### DNA and viral constructs

Parapinopsin in pcDNA3.1 was kindly provided by Akihisa Terakita (Osaka City University, Japan). EGFP-γ9 and mCherry-γ9 were previously generated (Karunarathne et al., 2012, 2013), and the PTx-resistant mutant Gα_OA_(C351I)-CFP was previously described (Azpiazu et al., 2006). Arrestin3-GFP was a gift from Ken Mackie (Indiana University Bloomington). Clathrin-mCherry was obtained from Addgene (#55019). RCaMP1.07 was a gift from Junichi Nakai (Saitama University, Japan).

mScarlet-γ9 was generated by PCR amplification of mScarlet using forward primer 5’-CCC AAG CTT ATG GTG AGC AAG GGC GAG GCA G-3’ (HindIII) and reverse primer 5’-CGC GGT ACC CTT GTA CAG CTC GTC CAT GCC G-3’ (KpnI). mCherry was excised from mCherry-γ9 (in pcDNA3.1) after HindIII and KpnI digest, and mScarlet was ligated in frame with γ9.

For *in vitro* characterization of purified protein, parapinopsin was subcloned into pFastbac vector (ThermoFisher) with an N-terminal FLAG tag and 10xHis tag. The construct was cloned using cut sites NcoI and XhoI through PCR amplification using forward primer 5’-CCA TCA CGC CAT GGA GAA CTT GAC C-3’ and reverse primer 5’-CCT CTA GAT GCA TGC TCG AGT CTA GCT CG-3’. PPO-Venus was generated by PCR amplification of PPO using forward primer 5’-GCG GAA TTC ATG GAG AAC TTG ACC TCG CTC GAC C-3’ (EcoR1) and reverse primer 5’-ATA GTT TAG CGG CCG CGC TCG GGG AGA CCT GCC CCG-3’ (NotI) and amplification of Venus using forward primer 5’-ATA AGA ATG CGG CCG CCA TGG TGA GCA AGG GCG AGG AGC-3’ (Not1 with extra base after the RE site to bring sequence into frame) and reverse primer 5’-GCT CTA GAT TAC TTG TAC AGC TCG TCC ATG CCG-3’ (XbaI site followed by a stop codon). Purified PCR products were ligated into pcDNA3.1 after vector digest with EcoR1 and XbaI.

Cre-dependent viral vectors were generated by amplifying PPO-Venus from pcDNA3.1. Resulting amplification products were restriction digested, then gel-purified and ligated into a Cre-dependent pAAV vector digested with the same enzymes used for the amplification product. The Cre-dependent viral vector using the CAG promoter to drive expression (kindly provided by Larry Zweifel, University of Washington) was digested with AgeI and AscI. PPO-Venus was amplified using the forward primer 5’-CGG ATC CAC TAG TCC AGA CCG GTG GAA TTC ATG G-3’ and the reverse primer 5’-GGC ACA GTC GAG GCG CGC CAG CGG GTT TAA ACG-3’, then digested with AgeI and AscI. The Cre-dependent viral vector using the EF1α promoter to drive expression (available from Addgene, plasmid # 27056) was digested with NheI and AscI. PPO-Venus was PCR amplified using the T7 promoter as a forward primer and the reverse primer 5’-GGC ACA GTC GAG GCG CGC CAG CGG GTT TAA ACG-3’, then ligated into the vector digested with NheI and AscI. All constructs were verified by Sanger sequencing.

AAVs were produced in the Hope Center Viral Vector Core. AAV5:CAG:DIO:PPO-Venus was used for all *in vitro* studies at a titer of 2*10^13^ vg/ml and AAV5:Ef1α:DIO:PPO-Venus for all *in vivo* studies at 3×10^13^ vg/ml. AAV5:Ef1α:DIO:eYFP was used for all control experiments at a titer of 5×10^12^ vg/ml.

### Antibodies

The following primary antibodies were used: mouse anti-synapsin-1 (Synaptic Systems cat. #106011, RRID:AB993033), guinea pig anti-tau (Synaptic Systems cat. # 314004, RRID:AB_A1547385), guinea pig anti-vGluT2 (Synaptic Systems cat. # 135404, RRID:AB_887884), and mouse anti-PSD-95 (NeuroMAb cat. # 75-028 RRID:AB_2292909). Secondary antibodies were both from Life Technologies: donkey anti-mouse Alexa Fluor 555 (cat. #31570) and goat anti-guinea pig Alexa Fluor 647 (cat. # 21450).

### Animals

All experiments were conducted in accordance with the National Institutes of Health guidelines and with approval from the Institutional Animal Care and Use Committee at Washington University School of Medicine and the Office of Animal Welfare at University of Washington. Postnatal 1-2 day male and female Sprague Dawley rats were used to prepare striatal cultures. 4-6 week old male *Avil^Cre^* mice (*Avil^tm2(cre)Fawa^;* Jackson labs #032536) were used for DRG cultures (da Silva et al., 2011; Zhou et al., 2010). Male and female *Vglut2-IRES-Cre* mice (*Slc17a6^tm2(Cre)Lowl^*; Jackson labs # 016963) were used for slice electrophysiology (Vong et al., 2011), and male and female *DAT-IRES-Cre* mice (*Slc6a3^tm1.1(Cre)Bkmn^*; Jackson labs # 006660) were used for behavioral experiments (Bäckman et al., 2006). All mice were group housed with *ad libitum* access to food and water before experiments and maintained on a 12hr light:dark cycle.

#### Stereotaxic viral injections and optic fiber surgeries

For slice electrophysiology, 5-6 week old *Vglut2-IRES-Cre* mice were anesthetized with isoflurane and secured on a stereotaxic frame. 1 μl of AAV5:Ef1α:DIO:PPO-Venus was injected unilaterally into the right VPm (AP: −1.5 mm, ML +1.8 mm, DV −3.6 mm) using a beveled tip Hamilton syringe. Viruses were injected at 75 nl/min and the needle was left in place for 10 minutes after infusion before being slowly withdrawn and suturing the skin.

In behavior experiments, 12-16 week old *DAT-IRES-Cre* (*Cre*^+^) mice or *Cre^-^* littermate controls were anesthetized with isoflurane and 0.4 μl (per hemisphere) of AAV5:Ef1α:DIO:PPO-Venus was injected bilaterally at 100 nl/min into the ventral tegmental area (VTA) (AP: −3.4 mm, ML: ±0.4 mm, DV: −4.4 mm). 220 μm optic fibers were placed bilaterally targeting either the VTA (coordinates same as above) or the nucleus accumbens core (NAc core) (AP: +1.4 mm, ML: +/− 1.5 mm, DV: −4.0 mm, 10° angle).

### *In vitro* protein characterization

#### Production of purified parapinopsin

The construct was grown in DH10Bac (ThermoFisher, 10361012) cells and transfected into Sf9 insect cells using Cellfectin II reagent (ThermoFisher, 10362100). Baculovirus was generated according to the manufacturer’s protocol. From this point onward all experiments were completed in the dark or under dim red light.

Sf9 cells were grown to a density of 2 million/mL at which time they were transduced with the baculovirus at a MOI=1.0. Protein was expressed for 48 hours in the dark. At 24 hours post-transduction, 10 µM 9-*cis-*retinal was added to the culture. Cells were harvested and spun down at 1000xg for 10 minutes. They were then resuspended in buffer containing 65mM NaCl, 50mM HEPES, 1mM EDTA (pH=7.4), with protease inhibitors partial thromboplastin time (PTT) and leupeptin hemisulfate (LS). The suspension was placed in a nitrogen cavitation chamber at 600 PSI for 30 minutes at 4°C. Upon release, lysed cells were centrifuged at 1000xg for 15 minutes followed by the supernatant being spun at 125,000xg for 35 minutes. The pellet was resuspended in buffer containing 50 mM NaCl, 50 mM HEPES, 5 mM MgCl_2_ (pH=7.4) with protease inhibitors (PTT/LS) and centrifuged at 125,000xg for 40 minutes. Resuspension of the pellet and centrifugation were repeated. Finally, the membranes were homogenized with a dounce and protein concentration was measured.

Membranes were diluted to 5mg/ml of total protein and solubilized in 200 mM NaCl, 20 mM HEPES, 5mM MgCl_2_, and 1% DDM (pH=7.4) at 4°C for 1 hour, at which point they were centrifuged at 125,000xg for 40 minutes. After centrifugation, the supernatant was applied to a Talon affinity column. The column was washed with 20 column volumes of wash buffer containing 50 mM NaCl, 20 mM HEPES, 5 mM MgCl_2_, 5 mM imidazole, and 0.1% DDM (pH=7.4). Protein was eluted with elution buffer containing 150 mM imidazole, 40 mM NaCl, 20 mM HEPES, 1mM MgCl_2_, 0.1% DDM (pH=7.4).

#### Spectroscopy

Absorption spectra of enriched protein samples were recorded on a SpectraMax 190 Microplate Reader (Molecular Devices) at 23°C. Cuvette was blanked with elution buffer. The protein sample was initially scanned in a dark state. A 1 kW halogen lamp (Phillips) with a shortpass filter (Edmund Optics) was used for blue light (470 nm) photoactivation of the protein and protein deactivation was achieved using filtered amber light (594nm). Absorbance measurements were plotted using Graphpad Prism.

### Cell culture and transfection

#### Real-time cAMP dynamics

Luminescence assays to monitor cAMP levels were performed as we described previously (Siuda et al., 2015). Briefly, HEK293 cells (ATCC, cat. # CRL-1573) were co-transfected with PPO-Venus and pGloSensor-22F (Promega E2301) plasmids using JetPrime reagent (Polyplus, cat. # 114-07). Two hours before the experiment, cells were incubated with 2% of the GloSensor reagent (Promega, cat. # E1290). Relative luminescence units were recorded using a SynergyMx microplate reader (Biotek). Adenylyl cyclase was activated with 1 μm forskolin, and cells were stimulated with constant blue LED light (465 nm, Plexon) for 60 seconds before being returned to the plate reader. Data, expressed as relative luminescence units, were normalized to the peak levels and plotted using GraphPad Prism.

#### Imaging assays of γ9 translocation

The HeLa cell line was obtained from ATCC and cultured in MEM supplemented with 10% dialyzed FBS (Atlanta Biologicals) and 1x pen-strep at 37°C and 5% CO_2_. Cells were cultured and transiently transfected in 35-mm glass bottom dishes (CellVis), using Lipofectamine 2000 (Invitrogen) according to the manufacturer’s protocol. Transfections used 2 μl of lipofectamine per dish along with the following DNA amounts:

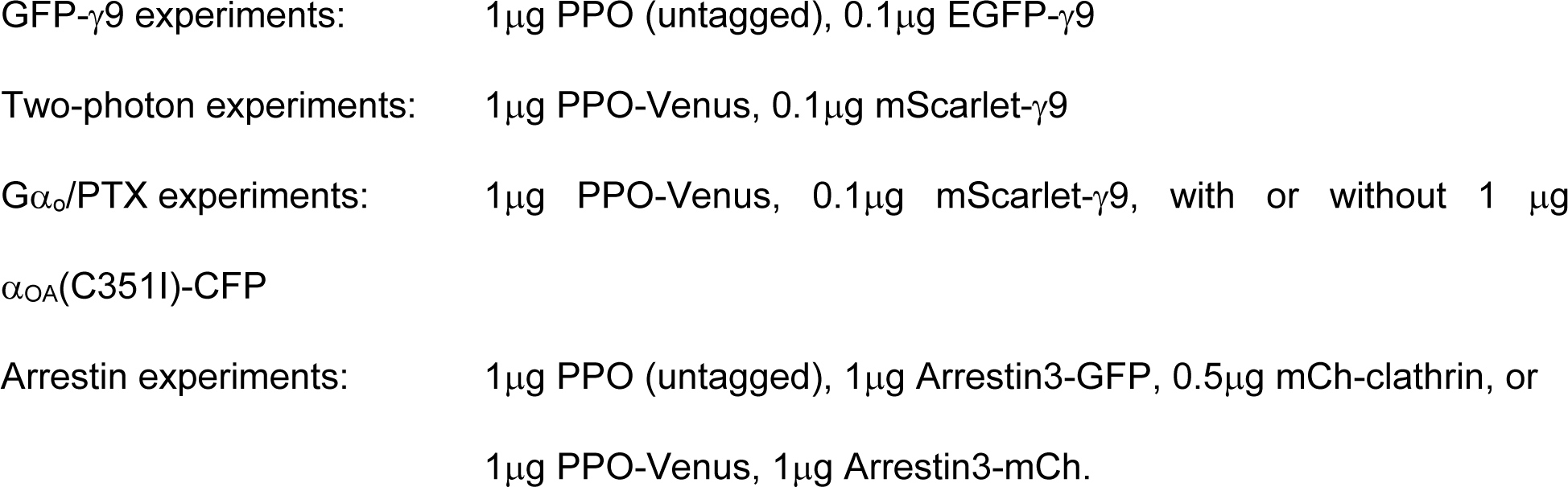

Cells were imaged 1-2 days after transfection.

#### Striatal neuron culture

Timed pregnant Sprague Dawley rats were purchased from Charles River. On postnatal day P1-P2, striatal cultures were prepared as detailed (Mao and Wang, 2001). Cells were plated at 50,000-100,000 cells per dish in the center wells of 35 mm dishes containing 10 mm glass bottom wells (CellVis). On the day before dissection, the dishes were coated with poly-D-lysine (PDL) by overnight incubation with 100 μg/ml PDL (Sigma P0899). Cells were cultured at 37°C and 5% CO2 in a 2:1 mix of NB:DMEM, where NB is neurobasal medium (Gibco) supplemented with 1x B27 (Gibco), 0.5mM glutamax (Gibco) and 1x pen-strep, and DMEM is DMEM/F12 medium (Gibco) supplemented with 10% FBS (Atlanta Biological), 1x B27, 10 g/l glucose, and 1x pen-strep. At 4DIV, half of the medium was exchanged for fresh NB+ containing Ara-C at a final concentration of 5 μM to prevent glial cell proliferation. Half the medium was exchanged for fresh NB+ every 7 days thereafter.

Cells were transfected using a modified version of the Lipofectamine 2000 transfection protocol, with increased reagent concentration but reduced transfection time, as follows. For each dish, 2 μl of lipofectamine and varying amounts of DNA (detailed below) were mixed in 100 μl of neurobasal medium and incubated at room temperature for 20 min. Conditioned medium was then removed from the dish of neurons, replaced with 100 μl DNA/lipofectamine mix, and placed back in the incubator. After 10 min, the transfection mix was removed from the dish and replaced with the conditioned medium.

PPO-Venus imaging: 0.5 μg PPO-Venus, 0.1 μg mScarlet-γ9, transfected 13DIV, imaged 14DIV

RCaMP imaging with PPO: 0.5μg PPO-Venus, 1 μg RCaMP1.07, transfected 7DIV, imaged 10DIV.

#### Dorsal root ganglia (DRG) neuron culture

DRG cultures were prepared from 4-6 week old *Avil^Cre^* mice as described previously (Siuda et al., 2015). Mice were anesthetized with isoflurane, spinal columns were removed, and DRG were dissected in HBSS + 10 mM HEPES (HBSS+H). Tissue was enzymatically digested with papain (45U, Worthington, cat. # LS003126) for 20 minutes at 37°C, rinsed in HBSS+H, followed by collagenase (1.5 mg/mL; Sigma, cat. #C6885) for 20 minutes at 37°C. Ganglia were washed and resuspended in culture media consisting of Neurobasal A (Gibco), 5% FBS (Life Technologies), 1x B27 (Gibco), 2 mM glutamax (Life Technologies) and 100 μg/mL penicillin/streptomycin (Life Technologies). Neurons were dissociated by mechanical trituration through glass pipettes, filtered with 40 μm filters and plated at a density of 10,000 cells/well onto poly-D-lysine and collagen coated 12 mm coverslips. Neurons were infected with 2*10^10^ vg/ml of Cre-dependent AAVs (AAV5:CAG:DIO:PPO-Venus or DIO:eYFP) 2 hours after plating. For experiments testing G-protein dependence, neurons were incubated overnight in 200 ng/ml pertussis toxin (List Biological Laboratories, cat. # 179A) before recordings.

### Histology

#### DRG cultures

One to 7 days after AAV infection, cultured DRG neurons were rinsed in PBS and fixed in 4% PFA/sucrose for 10 minutes on ice. After rinsing in PBS, neurons were blocked and permeabilized in PBS containing 3% donkey serum and 0.3% Triton X-100 for 15 minutes at room temperature. Cultures were incubated in primary antibodies (mouse anti-synapsin-1 and guinea pig anti-tau both at 1:2000 dilution) in PBS with 3% donkey serum overnight at 4°C. After washing, primary antibodies were labeled with donkey anti-mouse AF555 and goat anti-guinea pig AF647 secondary antibodies (both 1:2000 in 3% donkey serum) for 1 hour at room temperature. Coverslips were rinsed five times before being mounted on slides with Prolong Gold Antifade reagent with DAPI (Life Technologies, cat. # 36931). Neurons were imaged on a Leica TCS SPE confocal microscope with a 63x oil immersion objective (NA=1.4).

#### Thalamocortical projections

Two to 6 weeks after viral injections of Cre-dependent PPO into the VPM, mice were deeply anesthetized with ketamine/xylazine and transcardially perfused with PBS followed by 4% PFA. Brains were removed and cut at a 45° angle to the right of the midline to preserve thalamocortical axon projections in the plane of slicing. They were then post-fixed in PFA overnight, cryoprotected in 30% sucrose and frozen in OCT media. 50 μm thick thalamocortical slices were then cut parallel to the 45° plane and stored in PBS with 0.2% NaN_3_. Sections were rinsed thoroughly in PBS before being blocked and permeabilized in PBS with 2% donkey serum, 2% goat serum and 0.3% Triton X-100 for 1 hour at room temperature. Primary antibodies (mouse anti-PSD-95 (1:500) and guinea pig anti-vGluT2 (1:500)) were incubated with the sections overnight at 4°C in PBS with 2% each donkey and goat serum. Sections were washed 5 times and incubated with secondary antibodies (donkey anti-mouse AF555 (1:500) and goat anti-guinea pig AF647 (1:500)) for 1 hour at room temperature in PBS with 2% donkey and goat serum. Sections were washed 5 times again and mounted onto slides using Vectashield Hardset Antifade reagent with DAPI (Vector Labs, cat. # H-1400). Thalamocortical slice imaging was performed on an Andor Dragonfly microscope described below.

#### VTA dopamine neurons and projections

Following behavioral experiments, mice were deeply anesthetized with pentobarbital and transcardially perfused with PBS followed by 10% formalin. Heads were placed with optic fibers intact overnight in 10% formalin. Brains were extracted and post-fixed in formalin overnight, followed by cryoprotection in 30% sucrose. 30 μm thick slices containing either the VTA or NAc core were collected and mounted onto slides using Vectashield Hardset Antifade reagent with DAPI (Vector Labs, cat #H-1400) and imaged using an Olympus FV1000 confocal microscope (Olympus Scientific Solutions).

### Imaging assays

#### Live-cell confocal imaging

Imaging was performed using an Andor Dragonfly 500 spinning disk confocal system built on a Nikon Ti2 inverted fluorescence microscope equipped with Nikon Perfect Focus to actively prevent focal drift. The system is equipped with five solid state lasers (445 nm, 488 nm, 515 nm, 561 nm, and 640 nm), the emission can be collected along one of two paths containing emission filter wheels and either an EMCCD camera (Andor iXon Life) or a sCMOS camera (Andor Xyla). LED light from a multi-LED illumination system (CoolLED pE-4000) was directed through the back port of the microscope by an Andor Mosaic DMD, which enabled spatially patterned illumination at the sample. Blue or UV LED light was coupled into the excitation path with the 515nm and longer wavelength lasers using a dichroic mirror to enable simultaneous imaging and photoactivation. LED pulsing was controlled using the CoolLED touch pad and LED illumination was either applied at constant intensity, or in 10 ms pulses at 10Hz where specified.

HeLa cell imaging and RCaMP1.07 imaging in cultured neurons, used a 60x, 1.4NA oil immersion objective (Nikon MRD01605: CFI Plan Apo Lambda 60x Oil), the EMCCD camera, and single plane confocal imaging. Imaging of PPO-Venus distribution in transiently transfected striatal cultures used the same objective, but z-stacks at 210 nm steps were collected using the sCMOS camera and deconvolved using Andor Fusion software. An OkoLabs stage top incubator maintained the samples at 37°C, 5% CO_2_. HeLa cells and cultured neurons were imaged in their respective culture medium, supplemented with 10 μM 9-cis retinal.

#### Total internal reflection fluorescence (TIRF) imaging

TIRF imaging of PPO-induced arrestin translocation was performed on the Andor Dragonfly system described above, which is equipped for multi-color TIRF imaging. Images were captured using a 60x, 1.49 NA TIRF objective (Nikon: CFI Apochromat TIRF 60XC Oil) and an EMCCD camera (Andor iXon Life). Images were captured at 5 sec intervals for 10 min, and photoactivation with 460 nm LED light (10mW/mm^2^, 10 ms pulses, 10 Hz) was applied throughout the duration of the image sequence. Images of Arrestin3-GFP and mCh-clathrin were captured using 488 nm and 561 nm laser excitation, respectively. Images of PPO-Venus and Arrestin3-mCh were captured using 515 nm and 561 nm laser excitation, respectively.

#### Thalamocortical slice imaging

Brain slice imaging was performed using the Andor Dragonfly system described above. A 20x, 0.75NA objective (Nikon MRD00205) was used to identify regions of the barrel cortex for higher resolution imaging, which was performed using the 60x objective as above. Z-stack images were acquired using the sCMOS camera and deconvolved using Andor Fusion software. Laser excitation at 488nm, 561nm and 640nm was used to image PPO-Venus, Alexa Fluor 555 and 647 dyes, respectively.

#### LED-based deactivation during GFP imaging

Imaging was performed using an Andor Revolution spinning disk imaging system build on a Leica DMI6000B inverted microscope with adaptive focus control, with a Yokogawa CSU x1 spinning disk unit, and an Andor iXon EMCCD camera. The system is equipped with four solid state lasers at 445 nm, 488 nm, 514 nm, and 594 nm, combined through an acousto-optic tunable filter to enable rapid switching and control excitation power. An incubation chamber surrounding the entire microscope was maintained at 37°C, 5% CO_2_. Imaging was performed using a 63x, 1.4NA oil immersion objective (Leica 506187: HCX PL APO 63x). For GFP-γ9 imaging and simultaneous activation of PPO, a single confocal plane was imaged at a rate of one frame every 3 s, with 488 nm excitation through the spinning disk (average power ∼45 μW, which is typical for imaging GFP alone) and 300ms exposure time. PPO deactivation was achieved using a 595 nm LED (CoolLED pE-4000). The LED light was directed through the back port of the microscope by an Andor Mosaic DMD operating in “white mask” mode and ∼150 μW measured through the objective. The 488nm imaging laser and the 595 nm LED were coupled into the excitation path using a 562nm long pass dichroic mirror (Semrock Brightline FF556-SDi01). GFP emission was collected through the same dichroic mirror, a 488nm notch mirror, and a 525/30nm emission filter.

#### Multiphoton imaging

Multiphoton imaging was performed on an Olympus Fluoview FVMPE-RS multiphoton imaging system equipped with two wavelength-tunable Ti:Sapphire lasers (Spectra Physics, MaiTai and Insight lasers). HeLa cells transfected with PPO-Venus and mScarlet-γ9 were imaged at room temperature (22°C) in 35mm glass bottom dishes containing culture medium supplemented with 10 μM 9-cis retinal. A water immersion objective (Olympus XLPLN25XWMP2, 25x, 1.05NA) was immersed in the medium in the dish from above for imaging. Imaging and photostimulation were performed by laser scanning at a lateral resolution of 0.4792 μm/pixel. Imaging was performed with a scan speed of 20 μs/pixel, and each line was scanned 3 times and averaged. Two-photon excitation of mScarlet was performed using the Insight and tuned to 1080 nm. Two-photon photoactivation of PPO at each wavelength tested (700, 800, 900, 1000nm), was performed by tuning the wavelength of the MaiTai laser, adjusting the output power to an average of 10 mW measured at the back focal plane of the objective, and scanning the focused laser at a speed of 2 μs/pixel. The two lasers were combined into the same excitation path using a laser coupling dichroic mirror (Olympus LCDM690-1050). Emission was collected through a green-red filter cube (Olympus FV30-FGR) onto GaAsP photomultiplier tubes. In order to reset PPO into the off state after each image acquisition, the sample was illuminated from below through a condenser lens by constant amber LED light (ThorLabs M590L3) for at least 2 min.

#### Live-cell imaging analysis

Image sequences were first inspected manually for any lateral drift of the sample throughout the time course of the experiment. If necessary, the ImageJ TurboReg plugin was used to correct for lateral drift. Translocation of FP-γ9 from the plasma membrane to intracellular regions was quantified by manually selecting one cytosolic ROI per cell that excluded the plasma membrane and nucleus. The mean fluorescence intensity within the ROI was computed for each frame in a time series, using either ImageJ or Andor iQ software. An ROI selected in a nearby region of the sample lacking fluorescent cells was used for background subtraction. Since the fluorescence intensity varied among the transiently transfected cells, intracellular FP-γ9 intensities were normalized by the value at the first time point.

### Electrophysiology

#### Voltage gated calcium channel recordings

Calcium channel currents were recorded from cultured DRG neurons after 3-6 days *in vitro*. Neurons were transferred into choline external solution containing (in mM): 135 choline chloride, 10 HEPES, 10 glucose, 2 BaCl_2_, 4 MgCl_2_, pH adjusted to 7.3 with CsOH and 300-310 mOsm. 500 nM TTX (Abcam, cat. # ab120055) was included during recordings to block any residual sodium currents. Barium was used as the charge carrier for these recordings to avoid Ca^2+^ dependent changes in channel gating and physiology, and barium currents are referred to as calcium channel currents in the manuscript. GABA_B_Rs were stimulated with bath application of 50 μM R/S-baclofen (Abcam, cat. # ab120149), and 100 μM Cd^2+^ was routinely added at the end of an experiment to verify isolation of calcium channel currents. DRG neurons were visualized through a 40x objective using IR-DIC illumination to minimize light stimulation on an upright BX-51 microscope, and Venus/YFP-expressing neurons were identified using epifluorescence. Whole-cell patch-clamp recordings were made using fire-polished glass pipettes with resistances of 2-3 MΩ when filled with internal solution containing (in mM): 110 CsCl, 10 TEA chloride, 0.1 CaCl_2_, 10 EGTA, 10 HEPES, 4 MgATP, 0.4 Na_2_GTP, 10 Na_2_phosphocreatine, pH to 7.28 with CsOH, 290 mOsm. The majority of recordings were performed at 25°C. For testing the effects of PPO at physiological temperatures, a heated chamber (ALA Scientific Instruments, cat. # HCS) and temperature controller (ALA, cat. # HCT-10) were used to maintain the bath temperature at 32-34°C.

Whole-cell patch-clamp recordings were made with Patchmaster software controlling a HEKA EPC10 amplifier. Calcium channel currents were elicited by step depolarizations from −80 to 0 mV and monitored every 10 seconds, while leak currents were subtracted with a p/4 protocol of four 20 mV depolarizing test pulses after each sweep. IV curves were generated by step depolarizations from −80 to +60 mV in 10 mV increments. For pre-pulse protocols, neurons were depolarized to +80 mV for 50 ms, followed by a return to −80 mV for 10 ms before 50 ms test pulses to 0 mV. Currents were digitized at 20 kHz and filtered at 3 kHz. Whole-cell and pipette capacitance transients were minimized using the compensation circuits of the amplifier, and series resistance was compensated by 70% in all recordings. Only cells with stable R_s_ < 15 MΩ that did not change by more than 20% were included. Neurons were discarded from analysis if currents changed by more than 10% during baseline.

Photostimulation was delivered through the microscope objective from custom made LEDs coupled to the rear fluorescence port. Blue LED light (Thorlabs, 470L2) was collimated with an aspheric condenser lens (Thorlabs, cat. #ACL2520U-A) and directed into the microscope by a 505 nm longpass dichroic mirror, while collimated green (Thorlabs, 530L3) and amber (Thorlabs, 590L2) were coupled behind this mirror. Pulsed light was triggered by TTL pulses from a Master-9 signal generator (AMP Instruments) to the LED current controller (Thorlabs DC4104). All light intensities were calibrated using a photodiode (Thorlabs, S121C) and power meter (Thorlabs PM100D).

#### Acute slice preparation

Brain slices for electrophysiology recordings were prepared using a protective cutting and recovery method (Siuda et al., 2015; Ting et al., 2014). 2-6 weeks after viral injections, mice were deeply anesthetized with ketamine and xylazine and transcardially perfused with cold oxygenated NMDG-aCSF containing (in mM): 93 N-methyl-D-glucamine, 2.5 KCl, 1.25 NaH_2_PO_4_, 30 NaHCO_3_, 20 HEPES, 25 glucose, 5 ascorbic acid, 2 thiourea, 3 sodium pyruvate, 0.5 CaCl_2_, 5 MgCl_2_, pH=7.4, 300-310 mOsm. Brains were rapidly removed and blocked with a 45° cut to the right of the midline (Agmon and Connors, 1991; Crandall et al., 2017). The cut surface was then glued to a platform and embedded in 2% low-melt agarose (Sigma, A0676). 350 μm thick slices were cut using a Compresstome (Precisionary Instruments, cat. # VF-200-0Z) and transferred to a recovery chamber containing oxygenated NMDG-aCSF at 32°C for 10-12 minutes before moving to a holding chamber filled with room temperature oxygenated holding-aCSF containing (in mM): 92 NaCl, 2.5 KCl, 1.25 NaH_2_PO_4_, 30 NaHCO3, 20 HEPES, 25 glucose, 2 CaCl_2_, 2 MgCl_2_, pH=7.3, 300-310 mOsm. Slices were maintained in the dark at room temperature and allowed to recover >1 hour before recording.

#### Thalamocortical slice recordings

Slices were transferred to a heated recording chamber of an upright BX-51 microscope and perfused at a rate of 2 ml/minute with oxygenated aCSF containing (in mM): 124 NaCl, 2.5 KCl, 1.25 NaH_2_PO_4_, 24 NaHCO_3_, 5 HEPES, 12.5 glucose, 2 CaCl_2_, 1 MgCl_2_ (pH=7.3 and 300-310 mOSm). Slices were maintained at 32-34°C as described above. Neurons were visualized using IR-DIC microscopy and PPO-Venus+ fibers were imaged using LED epifluorescence through standard GFP filter cubes. Whole-cell patch-clamp recordings were made from visually identified neurons in layer IV of the barrel cortex using fire-polished glass pipettes with open tip resistances of 3-5 MΩ when filled with internal solution containing (in mM): 110 cesium gluconate, 8 tetraethylammonium chloride, 3 QX314 bromide, 1.1 EGTA, 0.1 CaCl_2_, 10 HEPES, 4 MgATP, 0.4 Na_2_GTP, 10 Na_2_phosphocreatine, pH to 7.27 with CsOH, 291 mOsm.

Recordings were performed using Patchmaster software controlling a HEKA EPC10. Neurons were voltage-clamped at −70 mV and electrically evoked thalamocortical currents were elicited by a bipolar stimulating electrode (FHC cat. #30202) placed in the VPm. Current pulses (0.2 ms, 50 ms interpulse intervals) was provided by a stimulus isolator (WPI, cat. #A365) and adjusted for each cell to produce ∼90% success rates. Excitatory postsynaptic currents were pharmacologically isolated using 10 μM bicuculline (Abcam, cat. # ab120108) and 100 μM picrotoxin (Abcam, cat. # ab120315) and elicited every 10 seconds. LED stimulation (10 Hz, 10 ms pulses, 10 mW/mm^2^) was delivered through the 40x objective above the recorded barreloid as described above. We confirmed the restricted photostimulation by illuminating PPO-Venus^+^ fibers within the 40x field of view using constant blue illumination (10 mW/mm^2^ for ∼5 minutes) and inspecting the photobleached area with the 4x objective. Series resistance values were not compensated and were continuously monitored using 5 mV hyperpolarizing test pulses every sweep. Neurons were excluded from analysis if R_s_ values changed more than 20% during recording. Data were sampled at 20 kHz, filtered at 3 kHz, and analyzed offline.

#### Electrophysiology analysis

Electrophysiology recordings were analyzed using Igor Pro software (Wavemetrics) with the NeuroMatic plug-in (Rothman and Silver, 2018) and custom-written macros. Calcium channel currents were normalized to membrane capacitance and plotted normalized to the average baseline amplitude. Neurons were discarded if baseline currents drifted by more than 10%. Percent inhibition was quantified as the maximum inhibition observed during optical or pharmacological stimulation relative to baseline amplitudes. EPSC amplitudes were taken as the average amplitude within 0.1 ms of the peak. Latencies and jitter were calculated from the onset of electrical stimulation to 10% of the peak rise-time. Series resistance changes were monitored by calculating the peak amplitudes of the capacitance transients during 5 mV hyperpolarizing test pulses. Recovery tau values were calculated from single-exponential fits of the normalized peak current amplitudes as they returned to baseline using NeuroMatic. Half maximal activation for IV curves were determined using Boltzmann sigmoidal non-linear regression with Graphpad Prism 8 software. Prism software was also used for all statistical analyses.

### Behavioral experiments

#### Operant behaviors

All behaviors were performed in a sound-attenuated room at least one week after habituation to handling and the holding room. Animals were trained on Pavlovian conditioning for 3 days, followed by fixed-ratio 1 (FR-1; 1 nose poke, one sucrose pellet) for 3 days, fixed ratio-3 (FR-3; 3 nose poke, one sucrose pellet) for 3 days and then progressive ratio testing. One week prior to Pavlovian conditioning, DAT-Cre+ or Cre-mice were food restricted down to ∼85% of free feeding body weight. Mice were also habituated to the reward (sucrose pellet) for 2 days prior to Pavlovian conditioning. Mice were trained to associate illumination of a house light (CS) with access to a sucrose pellet (US) within an operant box (Med-Associates, ENV-307A) under a variable interval 90 paradigm, where they would receive the pellet at a randomized interval of 30, 60 or 90 s separating consecutive trials. The house light would illuminate concurrently for 20s with reward delivery. A randomized intertrial interval of between 30-90 seconds separated consecutive trials. Pavlovian conditioning sessions lasted for 60 minutes, over which an average of 35 rewards were presented. Following conditioning for 3 days upon which animals consumed all the rewards presented in one session, mice underwent FR-1, where they were trained to perform 1 nose poke for 1 reward. Similar to Pavlovian conditioning, the house light would illuminate concurrently for 20s with reward delivery, following which, the animals would be able to proceed nose-poking. Upon optimum performance in FR-1, mice underwent FR-3 (3 nose-pokes for 1 reward) for 3 days, mice were counterbalanced to receive laser stimulation for photoinhibition at 10 Hz (473 nm wavelength, 10 ms pulse width, 5-8 mW of power; matched with optic fiber efficiency) for the entirety of the FR-3 session. Following photoinhibition during FR-3, mice were retrained on FR-3 without light stimulation and underwent progressive ratio (PR) testing, following the geometric progression, *nj* = 5*e ^j^*^/5^− 5, in which the criteria for rewards increased in an exponential manner (1, 2, 4, 6, 9, 12…) over the training period (Richardson and Roberts, 1996). Mice received laser stimulation for photoinhibition using the aforementioned parameters. Furthermore, during the photoinhibition experiments, mice were counterbalanced to receive laser stimulation during the entire PR session.

#### Conditioned place preference

*DAT-Cre^+^* or Cre^-^ mice were trained in an unbiased, three-compartment conditioning apparatus as described previously (Al-Hasani et al., 2015; McCall et al., 2015). In brief, we used a modified three-chamber CPP apparatus consisting of two square boxes (27 cm × 27 cm) that served as the conditioning chambers separated by a small center area that served as the passageway (5 cm wide × 8 cm long) between boxes. Boxes had 2.5 cm black-and-white vertical stripes or horizontal stripes and floors were covered with 500 ml of bedding on each side. The floor of the center area was smooth Plexiglas. Mice were habituated to handling, injection restraint and to the testing room for one week prior to testing. On day 1, mice were allowed free access to all 3 compartments of the conditioning apparatus for a 30-minute session. For 2 consecutive days following pre-test, mice received counterbalanced saline or cocaine (10 mg/kg, I.P) injections confined to either the left or right compartment at least 4 hours apart. Mice also received laser stimulation for photoinhibition at 10 Hz (473 nm wavelength, 10 ms pulse width, 5-8 mW of power; matched with optic fiber efficiency) on the cocaine-paired compartment for the entirety of the 30-minute session. Following conditioning, mice underwent a “post-test” day, where they were again allowed free access to all 3 compartments. Time spent in each chamber and total distance traveled for the 30-minute trial was measured using Ethovision 10 (Noldus Information Technologies, Leesburg, VA).

## REFERENCES

Agmon, A., and Connors, B.W. (1991). Thalamocortical responses of mouse somatosensory (barrel) cortex in vitro. Neuroscience 41, 365–379.

Airan, R.D., Thompson, K.R., Fenno, L.E., Bernstein, H., and Deisseroth, K. (2009). Temporally precise in vivo control of intracellular signalling. Nature 458, 1025–1029.

Al-Hasani, R., McCall, J.G., Shin, G., Gomez, A.M., Schmitz, G.P., Bernardi, J.M., Pyo, C.-O., Park, S. Il, Marcinkiewcz, C.M., Crowley, N.A., et al. (2015). Distinct subpopulations of nucleus accumbens dynorphin neurons drive aversion and reward. Neuron 87, 1063–1077.

Atasoy, D., and Sternson, S.M. (2018). Chemogenetic tools for causal cellular and neuronal biology. Physiol. Rev. 98, 391–418.

Azpiazu, I., Akgoz, M., Kalyanaraman, V., and Gautam, N. (2006). G protein βγ11 complex translocation is induced by Gi, Gq and Gs coupling receptors and is regulated by the α subunit type. Cell. Signal. 18, 1190–1200.

Bäckman, C.M., Malik, N., Zhang, Y., Shan, L., Grinberg, A., Hoffer, B.J., Westphal, H., and Tomac, A.C. (2006). Characterization of a mouse strain expressing Cre recombinase from the 3’ untranslated region of the dopamine transporter locus. Genesis 44, 383–390.

Bean, B.P. (1989). Neurotransmitter inhibition of neuronal calcium currents by changes in channel voltage dependence. Nature 340, 153–156.

Blackmer, T. (2001). G protein beta gamma subunit-mediated presynaptic inhibition: Regulation of exocytotic fusion downstream of Ca2+ entry. Science (80-.). 292, 293–297.

Bourinet, E., Soong, T.W., Stea, A., and Snutch, T.P. (1996). Determinants of the G protein-dependent opioid modulation of neuronal calcium channels. Proc. Natl. Acad. Sci. U. S. A. 93, 1486–1491.

Browning, K.N., Kalyuzhny, A.E., and Travagli, R.A. (2002). Opioid peptides inhibit excitatory but not inhibitory synaptic transmission in the rat dorsal motor nucleus of the vagus. J. Neurosci. 22, 2998–3004.

Burke, K.J., Keeshen, C.M., and Bender, K.J. (2018). Two forms of synaptic depression produced by differential neuromodulation of presynaptic calcium channels. Neuron 99, 969–984.e7.

Burnett, C.J., and Krashes, M.J. (2016). Resolving behavioral output via chemogenetic designer receptors exclusively activated by designer drugs. J. Neurosci. 36, 9268–9282.

Carrillo-Reid, L., Han, S., Yang, W., Akrouh, A., and Yuste, R. (2019). Controlling visually guided behavior by holographic recalling of cortical ensembles. Cell 178, 447–457.e5.

Copits, B.A., Pullen, M.Y., and Gereau, R.W. (2016). Spotlight on pain: optogenetic approaches for interrogating somatosensory circuits. Pain 157, 2424–2433.

Corre, J., van Zessen, R., Loureiro, M., Patriarchi, T., Tian, L., Pascoli, V., and Lüscher, C. (2018). Dopamine neurons projecting to medial shell of the nucleus accumbens drive heroin reinforcement. Elife 7, 1–22.

Crandall, S.R., Patrick, S.L., Cruikshank, S.J., and Connors, B.W. (2017). Infrabarrels are layer 6 circuit modules in the barrel cortex that link long-range inputs and outputs. Cell Rep. 21, 3065–3078.

Creed, M., Ntamati, N.R., Chandra, R., Lobo, M.K., and Lüscher, C. (2016). Convergence of reinforcing and anhedonic cocaine effects in the ventral pallidum. Neuron 92, 214–226.

Cruikshank, S.J., Lewis, T.J., and Connors, B.W. (2007). Synaptic basis for intense thalamocortical activation of feedforward inhibitory cells in neocortex. Nat. Neurosci. 10, 462– 468.

Currie, K.P.M. (2010). G protein inhibition of CaV2 calcium channels. Channels 4.

Dana, H., Mohar, B., Sun, Y., Narayan, S., Gordus, A., Hasseman, J.P., Tsegaye, G., Holt, G.T., Hu, A., Walpita, D., et al. (2016). Sensitive red protein calcium indicators for imaging neural activity. Elife 5, 1–24.

Davies, W.L., Hankins, M.W., and Foster, R.G. (2010). Vertebrate ancient opsin and melanopsin: divergent irradiance detectors. Photochem. Photobiol. Sci. 9, 1444–1457.

Dodt, E., and Meissl, H. (1982). The pineal and parietal organs of lower vertebrates. Experientia 38, 996–1000.

Dolphin, A.C., and Scott, R.H. (1987). Calcium channel currents and their inhibition by (-)- baclofen in rat sensory neurones: modulation by guanine nucleotides. J. Physiol. 386, 1–17.

Du, Y., Duc, N.M., Rasmussen, S.G.F., Hilger, D., Kubiak, X., Wang, L., Bohon, J., Kim, H.R., Wegrecki, M., Asuru, A., et al. (2019). Assembly of a GPCR-G protein complex. Cell 177, 1232–1242.e11.

Dunlap, K., and Fischbach, G.D. (1981). Neurotransmitters decrease the calcium conductance activated by depolarization of embryonic chick sensory neurones. J. Physiol. 317, 519–535.

Eichel, K., and von Zastrow, M. (2018). Subcellular organization of GPCR signaling. Trends Pharmacol. Sci. 39, 200–208.

Eichel, K., Jullié, D., and Von Zastrow, M. (2016). β-Arrestin drives MAP kinase signalling from clathrin-coated structures after GPCR dissociation. Nat. Cell Biol. 18, 303–310.

Eichel, K., Jullié, D., Barsi-Rhyne, B., Latorraca, N.R., Masureel, M., Sibarita, J.B., Dror, R.O., and Von Zastrow, M. (2018). Catalytic activation of β-Arrestin by GPCRs. Nature 557, 381–386.

Eickelbeck, D., Rudack, T., Tennigkeit, S.A., Surdin, T., Karapinar, R., Schwitalla, J.C., Mücher, B., Shulman, M., Scherlo, M., Althoff, P., et al. (2019). Lamprey parapinopsin (“UVLamP”): a bistable UV-sensitive optogenetic switch for ultrafast control of GPCR pathways. ChemBioChem 1–7.

Ernst, O.P., Lodowski, D.T., Elstner, M., Hegemann, P., and Brown, L.S. (2014). Microbial and animal rhodopsins.

Gunaydin, L.A., Grosenick, L., Finkelstein, J.C., Kauvar, I. V, Fenno, L.E., Adhikari, A., Lammel, S., Mirzabekov, J.J., Airan, R.D., Zalocusky, K.A., et al. (2014). Natural neural projection dynamics underlying social behavior. Cell 157, 1535–1551.

Heinke, B., Gingl, E., and Sandkühler, J. (2011). Multiple targets of μ-opioid receptor-mediated presynaptic inhibition at primary afferent Aδ- and C-fibers. J. Neurosci. 31, 1313–1322.

Herlitze, S., Garcia, D.E., Mackie, K., Hille, B., Scheuer, T., and Catterall, W.A. (1996). Modulation of Ca2+ channels by G-protein beta gamma subunits. Nature 380, 258–262.

Hodos, W. (1961). Progressive ratio as a measure of reward strength. Science 134, 943–944.

Hunt, T.W., Carroll, R.C., and Peralta, E.G. (1994). Heterotrimeric G proteins containing G alpha i3 regulate multiple effector enzymes in the same cell. Activation of phospholipases C and A2 and inhibition of adenylyl cyclase. J. Biol. Chem. 269, 29565–29570.

Ikeda, S.R. (1996). Voltage-dependent modulation of N-type calcium channels by G-protein beta gamma subunits. Nature 380, 255–258.

Inoue, M., Takeuchi, A., Manita, S., Horigane, S. ichiro, Sakamoto, M., Kawakami, R., Yamaguchi, K., Otomo, K., Yokoyama, H., Kim, R., et al. (2019). Rational engineering of XCaMPs, a multicolor GECI suite for in vivo imaging of complex brain circuit dynamics. Cell 177, 1346–1360.e24.

Johnson, K.A., and Lovinger, D.M. (2016). Presynaptic G protein-coupled receptors: Gatekeepers of addiction? Front. Cell. Neurosci. 10, 1–22.

Karunarathne, W.K.A., O’Neill, P.R., Martinez-Espinosa, P.L., Kalyanaraman, V., and Gautam, N. (2012). All G protein βγ complexes are capable of translocation on receptor activation.

Biochem. Biophys. Res. Commun. 421, 605–611.

Karunarathne, W.K.A., Giri, L., Kalyanaraman, V., and Gautam, N. (2013). Optically triggering spatiotemporally confined GPCR activity in a cell and programming neurite initiation and extension. Proc. Natl. Acad. Sci. U. S. A. 110, 1565–1574.

Katada, T., and Ui, M. (1982). Direct modification of the membrane adenylate cyclase system by islet-activating protein due to ADP-ribosylation of a membrane protein (rat C6 glioma cel/3-adrenergic receptor/NAD/guanine nucleotide regulatory protein). Biochemistry 79, 3129–3133.

Kato, H.E., Zhang, Y., Hu, H., Suomivuori, C.M., Kadji, F.M.N., Aoki, J., Krishna Kumar, K., Fonseca, R., Hilger, D., Huang, W., et al. (2019). Conformational transitions of a neurotensin receptor 1–Gi1 complex. Nature 572, 80–85.

Kawano-Yamashita, E., Koyanagi, M., Shichida, Y., Oishi, T., Tamotsu, S., and Terakita, A. (2011). Beta-arrestin functionally regulates the non-bleaching pigment parapinopsin in lamprey pineal. PLoS One 6, 4–5.

Kawano-Yamashita, E., Koyanagi, M., Wada, S., Tsukamoto, H., Nagata, T., and Terakita, A. (2015). Activation of transducin by bistable pigment parapinopsin in the pineal organ of lower vertebrates. PLoS One 10, 1–13.

Kim, C.K., Adhikari, A., and Deisseroth, K. (2017). Integration of optogenetics with complementary methodologies in systems neuroscience. Nat. Rev. Neurosci. 18, 222–235.

Kleinlogel, S. (2016). Optogenetic user’s guide to Opto-GPCRs. Front. Biosci. (Landmark Ed. 21, 794–805.

Kouyama, T., and Murakami, M. (2010). Structural divergence and functional versatility of the rhodopsin superfamily. Photochem. Photobiol. Sci. 9, 1458–1465.

Koyanagi, M., and Terakita, A. (2014). Diversity of animal opsin-based pigments and their optogenetic potential. Biochim. Biophys. Acta - Bioenerg. 1837, 710–716.

Koyanagi, M., Kawano, E., Kinugawa, Y., Oishi, T., Shichida, Y., Tamotsu, S., and Terakita, A. (2004). Bistable UV pigment in the lamprey pineal. Proc. Natl. Acad. Sci. 101, 6687–6691.

Koyanagi, M., Kawano-Yamashita, E., Wada, S., and Terakita, A. (2017). Vertebrate bistable pigment parapinopsin: Implications for emergence of visual signaling and neofunctionalization of non-visual pigment. Front. Ecol. Evol. 5, 1–7.

Kreitzer, A.C., and Malenka, R.C. (2007). Endocannabinoid-mediated rescue of striatal LTD and motor deficits in Parkinson’s disease models. Nature 445, 643–647.

Li, P., Rial, D., Canas, P.M., Yoo, J., Li, W., Zhou, X., Wang, Y., van Westen, G.J.P., Payen, M., Augusto, E., et al. (2015). Optogenetic activation of intracellular adenosine A2A receptor signaling in the hippocampus is sufficient to trigger CREB phosphorylation and impair memory. Mol. Psychiatry 20, 1–11.

Lin, J.Y., Sann, S.B., Zhou, K., Nabavi, S., Proulx, C.D., Malinow, R., Jin, Y., and Tsien, R.Y. (2013). Optogenetic inhibition of synaptic release with chromophore-assisted light inactivation (CALI). Neuron 79, 241–253.

Liu, Q., Sinnen, B.L., Boxer, E.E., Schneider, M.W., Grybko, M.J., Buchta, W.C., Gibson, E.S., Wysoczynski, C.L., Ford, C.P., Gottschalk, A., et al. (2019). A Photoactivatable Botulinum Neurotoxin for Inducible Control of Neurotransmission. Neuron 101, 863–875.e6.

Lobingier, B.T., and von Zastrow, M. (2019). When trafficking and signaling mix: How subcellular location shapes G protein-coupled receptor activation of heterotrimeric G proteins. Traffic 20, 130–136.

Lüscher, C., and Slesinger, P. a (2010). Emerging roles for G protein-gated inwardly rectifying potassium (GIRK) channels in health and disease. Nat. Rev. Neurosci. 11, 301–315.

Mahn, M., Prigge, M., Ron, S., Levy, R., and Yizhar, O. (2016). Biophysical constraints of optogenetic inhibition at presynaptic terminals. Nat. Neurosci. 19, 554–556.

Mahn, M., Gibor, L., Patil, P., Cohen-Kashi Malina, K., Oring, S., Printz, Y., Levy, R., Lampl, I., and Yizhar, O. (2018). High-efficiency optogenetic silencing with soma-targeted anion-conducting channelrhodopsins. Nat. Commun. 9.

Mahn, M., Saraf-Sinik, I., Patil, P., Pulin, M., Bruentgens, F., Bitton, E., Palgi, S., Gat, A., Dine, J., Wietek, J., et al. (2020). Optogenetic silencing of neurotransmitter release with a naturally occurring invertebrate rhodopsin. Submitted.

Mao, L., and Wang, J.Q. (2001). Upregulation of preprodynorphin and preproenkephalin mRNA expression by selective activation of group I metabotropic glutamate receptors in characterized primary cultures of rat striatal neurons. Mol. Brain Res. 86, 125–137.

Marcott, P.F., Gong, S., Donthamsetti, P., Grinnell, S.G., Nelson, M.N., Newman, A.H., Birnbaumer, L., Martemyanov, K.A., Javitch, J.A., and Ford, C.P. (2018). Regional Heterogeneity of D2-Receptor Signaling in the Dorsal Striatum and Nucleus Accumbens. Neuron 98, 575–587.e4.

Marshall, J., Carleton, K.L., and Cronin, T. (2015). Colour vision in marine organisms. Curr. Opin. Neurobiol. 34, 86–94.

Marshel, J.H., Kim, Y.S., Machado, T.A., Quirin, S., Benson, B., Kadmon, J., Raja, C., Chibukhchyan, A., Ramakrishnan, C., Inoue, M., et al. (2019). Cortical layer-specific critical dynamics triggering perception. Science (80-.). 365.

Mattis, J., Tye, K.M., Ferenczi, E.A., Ramakrishnan, C., O’Shea, D.J., Prakash, R., Gunaydin, L.A., Hyun, M., Fenno, L.E., Gradinaru, V., et al. (2011). Principles for applying optogenetic tools derived from direct comparative analysis of microbial opsins. Nat. Methods 9, 159–172.

McCall, J.G., Al-Hasani, R., Siuda, E.R., Hong, D.Y., Norris, A.J., Ford, C.P., and Bruchas, M.R. (2015). CRH engagement of the locus coeruleus noradrenergic system mediates stress-induced anxiety. Neuron 87, 605–620.

Messier, J.E., Chen, H., Cai, Z.L., and Xue, M. (2018). Targeting light-gated chloride channels to neuronal somatodendritic domain reduces their excitatory effect in the axon. Elife 7, 1–21.

Morales, M., and Margolis, E.B. (2017). Ventral tegmental area: Cellular heterogeneity, connectivity and behaviour. Nat. Rev. Neurosci. 18, 73–85.

Morita, Y., Tabata, M., Uchida, K., and Samejima, M. (1992). Pineal-dependent locomotor activity of lamprey, Lampetra japonica, measured in relation to LD cycle and circadian rhythmicity. J. Comp. Physiol. A 171, 555–562.

Nanou, E., and Catterall, W.A. (2018). Calcium Channels, Synaptic Plasticity, and Neuropsychiatric Disease. Neuron 98, 466–481.

Nuber, S., Zabel, U., Lorenz, K., Nuber, A., Milligan, G., Tobin, A.B., Lohse, M.J., and Hoffmann, C. (2016). B-Arrestin biosensors reveal a rapid, receptor-dependent activation/deactivation cycle. Nature 531, 1–13.

O’Neill, P.R., Karunarathne, W.K.A., Kalyanaraman, V., Silvius, J.R., and Gautama, N. (2012). G-protein signaling leverages subunit-dependent membrane affinity to differentially control βγ translocation to intracellular membranes. Proc. Natl. Acad. Sci. U. S. A. 109.

O’Neill, P.R., Castillo-Badillo, J.A., Meshik, X., Kalyanaraman, V., Melgarejo, K., and Gautam, N. (2018). Membrane Flow Drives an Adhesion-Independent Amoeboid Cell Migration Mode. Dev. Cell 46, 9–22.e4.

Oh, E., Maejima, T., Liu, C., Deneris, E., and Herlitze, S. (2010). Substitution of 5-HT1A receptor signaling by a light-activated G protein-coupled receptor. J. Biol. Chem. 285, 30825– 30836.

Ohkura, M., Sasaki, T., Kobayashi, C., Ikegaya, Y., and Nakai, J. (2012). An improved genetically encoded red fluorescent Ca2+ indicator for detecting optically evoked action potentials. PLoS One 7.

Packer, A.M., Russell, L.E., Dalgleish, H.W.P., and Häusser, M. (2014). Simultaneous all-optical manipulation and recording of neural circuit activity with cellular resolution in vivo. Nat. Methods 12, 140–146.

Parker, K.E., Pedersen, C.E., Gomez, A.M., Spangler, S.M., Walicki, M.C., Feng, S.Y., Stewart, S.L., Otis, J.M., Al-Hasani, R., McCall, J.G., et al. (2019). A Paranigral VTA Nociceptin Circuit that Constrains Motivation for Reward. Cell 178, 653–671.e19.

Raimondo, J. V, Kay, L., Ellender, T.J., and Akerman, C.J. (2012). Optogenetic silencing strategies differ in their effects on inhibitory synaptic transmission. Nat. Neurosci. 15, 1102– 1104.

Richardson, N.R., and Roberts, D.C. (1996). Progressive ratio schedules in drug self-administration studies in rats: a method to evaluate reinforcing efficacy. J. Neurosci. Methods 66, 1–11.

Rickgauer, J.P., Deisseroth, K., and Tank, D.W. (2014). Simultaneous cellular-resolution optical perturbation and imaging of place cell firing fields. Nat. Neurosci. 17, 1816–1824.

Rost, B.R., Schneider-Warme, F., Schmitz, D., and Hegemann, P. (2017). Optogenetic Tools for Subcellular Applications in Neuroscience. Neuron 96, 572–603.

Roth, B.L. (2016). DREADDs for Neuroscientists. Neuron 89, 683–694.

Rothman, J.S., and Silver, R.A. (2018). NeuroMatic: An Integrated Open-Source Software Toolkit for Acquisition, Analysis and Simulation of Electrophysiological Data. Front. Neuroinform. 12, 1–21.

Scott, R.H., and Dolphin, A.C. (1987). Activation of a G protein promotes agonist responses to calcium channel ligands. Nature 330, 760–762.

da Silva, S., Hasegawa, H., Scott, A., Zhou, X., Wagner, A.K., Han, B.-X., and Wang, F. (2011). Proper formation of whisker barrelettes requires periphery-derived Smad4-dependent TGF-beta signaling. Proc. Natl. Acad. Sci. U. S. A. 108, 3395–3400.

Siuda, E.R., Copits, B.A., Schmidt, M.J., Baird, M.A., Al-Hasani, R., Planer, W.J., Funderburk, S.C., McCall, J.G., Gereau IV, R.W., and Bruchas, M.R. (2015). Spatiotemporal control of opioid signaling and behavior. Neuron 86, 923–935.

Stachniak, T.J., Ghosh, A., and Sternson, S.M. (2014). Chemogenetic Synaptic Silencing of Neural Circuits Localizes a Hypothalamus→Midbrain Pathway for Feeding Behavior. Neuron 82, 797–808.

Stujenske, J.M., Spellman, T., and Gordon, J.A. (2015). Modeling the Spatiotemporal Dynamics of Light and Heat Propagation for InVivo Optogenetics. Cell Rep. 12, 525–534.

Takahashi, T., Kajikawa, Y., and Tsujimoto, T. (1998). G-protein-coupled modulation of presynaptic calcium currents and transmitter release by a GABA(B) receptor. J. Neurosci. 18, 3138–3146.

Tejeda, H.A., Wu, J., Kornspun, A.R., Pignatelli, M., Kashtelyan, V., Krashes, M.J., Lowell, B.B., Carlezon, W.A., and Bonci, A. (2017). Pathway- and Cell-Specific Kappa-Opioid Receptor Modulation of Excitation-Inhibition Balance Differentially Gates D1 and D2 Accumbens Neuron Activity. Neuron 93, 147–163.

Terakita, A., Nagata, T., Sugihara, T., and Koyanagi, M. (2015). Optogenetic Potentials of Diverse Animal Opsins. In Optogenetics: Light-Sensing Proteins and Their Applications, H. Yawo, H. Kandori, and A. Koizumi, eds. (Tokyo: Springer Japan), pp. 77–88.

Ting, J.T., Daigle, T.L., Chen, Q., and Feng, G. (2014). Methods and Protocols. 1183, 1–629.

Touhara, K.K., and Mackinnon, R. (2018). Molecular basis of signaling specificity between GIRK channels and GPCRs. Elife 7, 1–23.

Tye, K.M., Mirzabekov, J.J., Warden, M.R., Ferenczi, E.A., Tsai, H.C., Finkelstein, J., Kim, S.Y., Adhikari, A., Thompson, K.R., Andalman, A.S., et al. (2013). Dopamine neurons modulate neural encoding and expression of depression-related behaviour. Nature 493, 537–541.

Uchida, K., and Morita, Y. (1994). Spectral sensitivity and mechanism of interaction between inhibitory and excitatory responses of photosensory pineal neurons. Pflügers Arch. Eur. J. Physiol. 427, 373–377.

Vong, L., Ye, C., Yang, Z., Choi, B., Chua, S., and Lowell, B.B. (2011). Leptin Action on GABAergic Neurons Prevents Obesity and Reduces Inhibitory Tone to POMC Neurons. Neuron 71, 142–154.

Wiegert, J.S., Mahn, M., Prigge, M., Printz, Y., and Yizhar, O. (2017). Silencing Neurons: Tools, Applications, and Experimental Constraints. Neuron 95, 504–529.

Wilden, U., Wüst, E., Weyand, I., and Kühn, H. (1986). Rapid affinity purification of retinal arrestin (48 kDa protein) via its light-dependent binding to phosphorylated rhodopsin. FEBS Lett. 207, 292–295.

Wingler, L.M., Elgeti, M., Hilger, D., Latorraca, N.R., Lerch, M.T., Staus, D.P., Dror, R.O., Kobilka, B.K., Hubbell, W.L., and Lefkowitz, R.J. (2019). Angiotensin Analogs with Divergent Bias Stabilize Distinct Receptor Conformations. Cell 176, 468–478.e11.

Wise, A., Watson-Koken, M.A., Rees, S., Lee, M., and Milligan, G. (1997). Interactions of the α2A-adrenoceptor with multiple Gi-family G-proteins: Studies with pertussis toxin-resistant G-protein mutants. Biochem. J. 321, 721–728.

van Wyk, M., Pielecka-Fortuna, J., Löwel, S., and Kleinlogel, S. (2015). Restoring the ON switch in blind retinas: opto-mGluR6, a next-generation, cell-tailored optogenetic tool. PLoS Biol. 13, e1002143.

Yang, W., Carrillo-Reid, L., Bando, Y., Peterka, D.S., and Yuste, R. (2018). Simultaneous two-photon imaging and two-photon optogenetics of cortical circuits in three dimensions. Elife 7, 1– 21.

Zamponi, G.W., and Currie, K.P.M. (2013). Regulation of Ca(V)2 calcium channels by G protein coupled receptors. Biochim. Biophys. Acta 1828, 1629–1643.

Zhou, X., Wang, L., Hasegawa, H., Amin, P., Han, B.-X., Kaneko, S., He, Y., and Wang, F. (2010). Deletion of PIK3C3/Vps34 in sensory neurons causes rapid neurodegeneration by disrupting the endosomal but not the autophagic pathway. Proc. Natl. Acad. Sci. U. S. A. 107, 9424–9429.

Zucker, R.S., and Regehr, W.G. (2002). Short-Term Synaptic Plasticity. Annu. Rev. Physiol. 64, 355–405.

Zurawski, Z., Gray, A.D.T., Brady, L.J., Page, B., Church, E., Harris, N.A., Dohn, M.R., Yim, Y.Y., Hyde, K., Mortlock, D.P., et al. (2019a). Disabling the Gβγ-SNARE interaction disrupts GPCR-mediated presynaptic inhibition, leading to physiological and behavioral phenotypes. Sci. Signal. 12.

Zurawski, Z., Yim, Y.Y., Alford, S., and Hamm, H.E. (2019b). The expanding roles and mechanisms of G protein–mediated presynaptic inhibition. J. Biol. Chem. 294, 1661–1670.

