## Supplemental Figures for "A photoswitchable GPCR-based opsin for presynaptic silencing"

**Supplemental Figure 1**

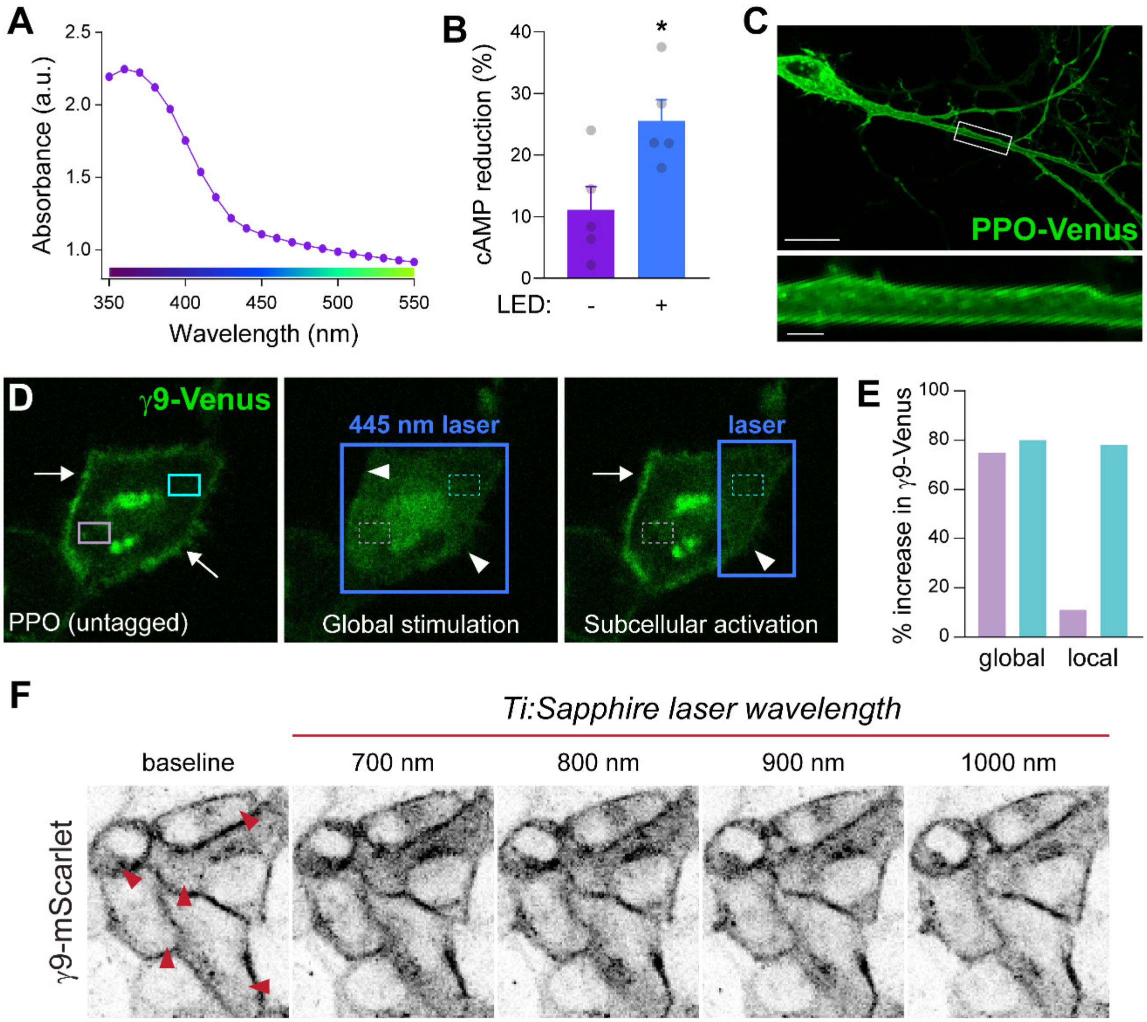


**Supplemental Figure 1. Trafficking and dynamics of PPO activation *in vitro* (related to Figure 1)**

(A) Absorbance values of purified PPO proteins in the dark state when illuminated with 350-550 nm light.

(B) Quantification of the decrease in cAMP luminescence in PPO-expressing HEK cells after 1 minute of constant illumination with 465 nm light or no LED stimulation. Adenylyl cyclase activity was stimulated with 1 μM forskolin and values were normalized to cAMP luminescence before LED stimulation.

(C) Confocal image of PPO-Venus fluorescence 1 day after transfection in cultured rat striatal neurons. PPO exhibited a high level of membrane expression and was effectively trafficked into distal dendrites. Inset depicts membrane localization in a secondary dendrite. The scale bars are 1 and 10 μm for the cropped and full field of view, respectively.

(D) Images of γ9-Venus translocation in response to cellular and subcellular activation by a 445 nm laser in HeLa cells. Arrows point to plasma membrane localized γ9-Venus fluorescence, which translocates to local intracellular membranes during photostimulation (marked by arrow heads). Translocation was quantified in the violet and teal regions of interest (ROIs) marked by dashed outlines.

(E) Quantification of γ9-Venus translocation during global stimulation of the cell or local activation in the violet or teal ROIs in panel C.

(F) Images of γ9-mScarlet translocation in HeLa cells when stimulated with different wavelengths from a tunable Ti:Sapphire laser. γ9-mScarlet was imaged with a second laser tuned to 1080 nm. Red arrowheads highlight areas of translocation.

### Supplemental Figure 2

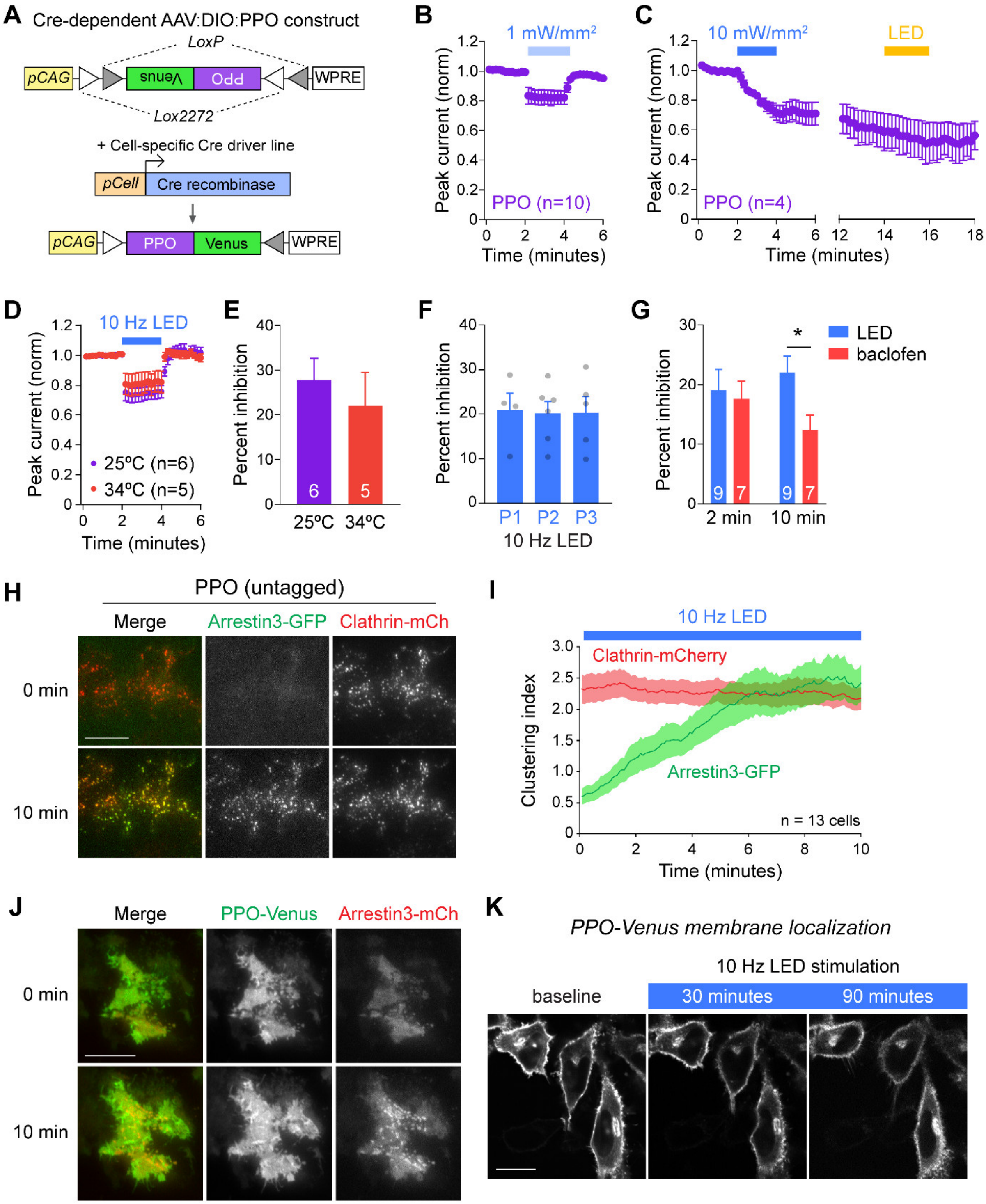

**Supplemental Figure 2. Characterization of PPO $Ca^{2+}$ channel inhibition, desensitization, and internalization (related to Figure 2).**

(A) Cartoon of the Cre-dependent PPO adeno-associated viral (AAV) construct. This double-floxed inverted (DIO) PPO is in an inverted orientation and flanked by non-homologous recombination sites (*LoxP* and *Lox2272*). Co-expression with Cre recombinase flips the construct and excises these sites, resulting in proper orientation of the PPO-Venus transgene, with expression driven by the upstream CAG promoter.

(B) Plot of normalized Ca2+ channel currents in DRG neurons. Blue bar indicates constant photostimulation with 470 nm light at 1 $mW/mm^2$. Note that these currents recovered rapidly after LED was turned off.

(C) Plot of normalized $Ca^{2+}$ channel currents in response to constant illumination with higher intensity light at 10 $mW/mm^2$. These currents did not spontaneously recover after stimulation and inhibition could not be switched off with amber light (590 nm, 1 $mW/mm^2$).

(D) Plot of normalized $Ca^{2+}$ channel responses to pulsed blue light (10 Hz, 10 ms, 10 $mW/mm^2$) at ambient (25°C, purple) and physiological temperatures (34°C, red).

(E) Summary graph of $Ca^{2+}$ channel inhibition at 25°C and 34°C showing no effects of temperature on the magnitude of inhibition.

(F) Summary graph of the percent inhibition of $Ca^{2+}$ channel currents to three repeated LED pulses (P1-P3) depicted in Figure 2F.

(G) Summary graphs of $Ca^{2+}$ channel inhibition at 2 and 10 minutes after stimulation of PPO with 10 Hz LED light (blue bars) or endogenous $GABA_B$Rs with baclofen (coral) as shown in Figure 2G $^*p<0.05$

(H) Live-cell total internal reflection fluorescence (TIRF) microscopy images of GFP-tagged arrestin3 (green) and mCherry-tagged clathrin (red) within ~100 nm of the plasma membrane in HeLa cells co-expressing untagged PPO. Stimulating PPO with 10 Hz blue light-induced clustering of arrestin3-GFP at endocytic zones marked by clathrin-mCherry. Scale bar represents 10 μm.

(I) Quantification of arrestin-GFP and clathrin-mCherry clustering during PPO stimulation.

(J) Live-cell TIRF microscopy images of arrestin3-mCherry (red) and PPO-Venus (green) in HeLa cells. LED stimulation caused arrestin3 activation and clustering but did not induce PPO-Venus trafficking to these endocytic zones. Scale bar represents 10 μm.

(K) Live confocal imaging of PPO-Venus trafficking in HeLa cells following prolonged optical stimulation. We did not observe changes in membrane localization or endocytosis of PPO even after 90 minutes of 10 Hz LED stimulation. Images have been normalized to adjust for photobleaching. Scale bar represents 10 μm.

**Supplemental Figure 3**

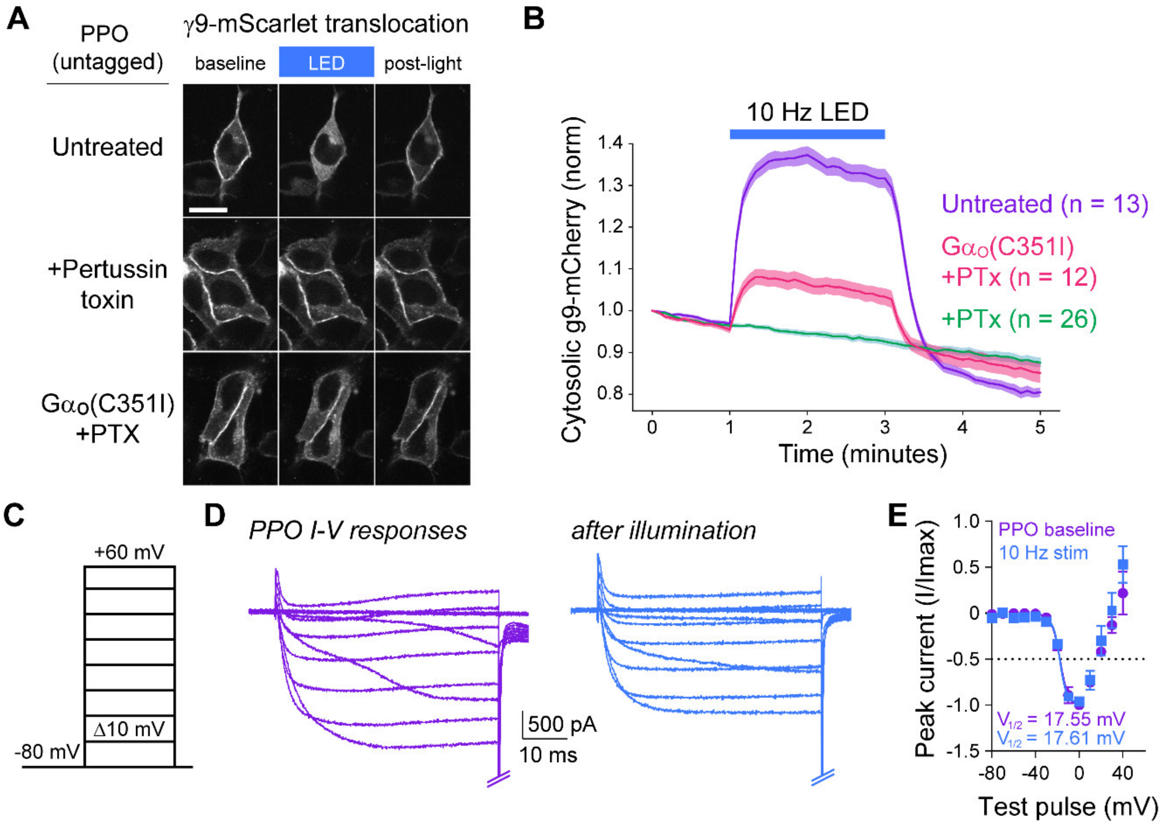


**Supplemental Figure 3. PPO can also couple to Ga$_o$ proteins but does not alter Ca$^{2+}$ channel gating (related to Figure 3).**

(A) Live-cell confocal images of γ9-mScarlet translocation in HeLa cells co-expressing PPO. LED stimulation induces prominent translocation of γ9-mScarlet from the plasma membrane which is completely blocked by the G$\alpha_{i/o}$ inhibitor pertussis toxin (PTx). Co-expression of a PTx-resistant G$\alpha_o$ mutant (C351I) rescues this translocation, demonstrating that PPO can also couple to G$\alpha_o$ proteins. Scale bar represents 10 μm.

(B) Quantification of γ9-mScarlet translocation following LED stimulation of PPO. Translocation is completely blocked by PTx and partially rescued by PTx-resistant G$\alpha_o$(C351) mutants.

(C) Diagram of voltage steps from -80 to +60 mV in 10 mV increments that were used to generate I-V plots.

(D) Overlaid $Ca^{2+}$ channel current traces elicited by 50 ms step depolarizations shown in panel C. Inward currents were activated around -30 mV and peaked at 0 mV. Baseline PPO responses are shown in purple and blue traces depict I-V relationships generated after stimulation with blue LED light.

(E) Current-voltage (I-V) plot of normalized currents before (purple) and after LED stimulation (blue). Despite decreasing $Ca^{2+}$ channel amplitudes, PPO did not affect voltage-dependent gating. Activation curves were fit using Boltzmann sigmoidal non-linear regression, which revealed no change in half-maximal activation voltage ($V_{1/2}$).

**Supplemental Figure 4**

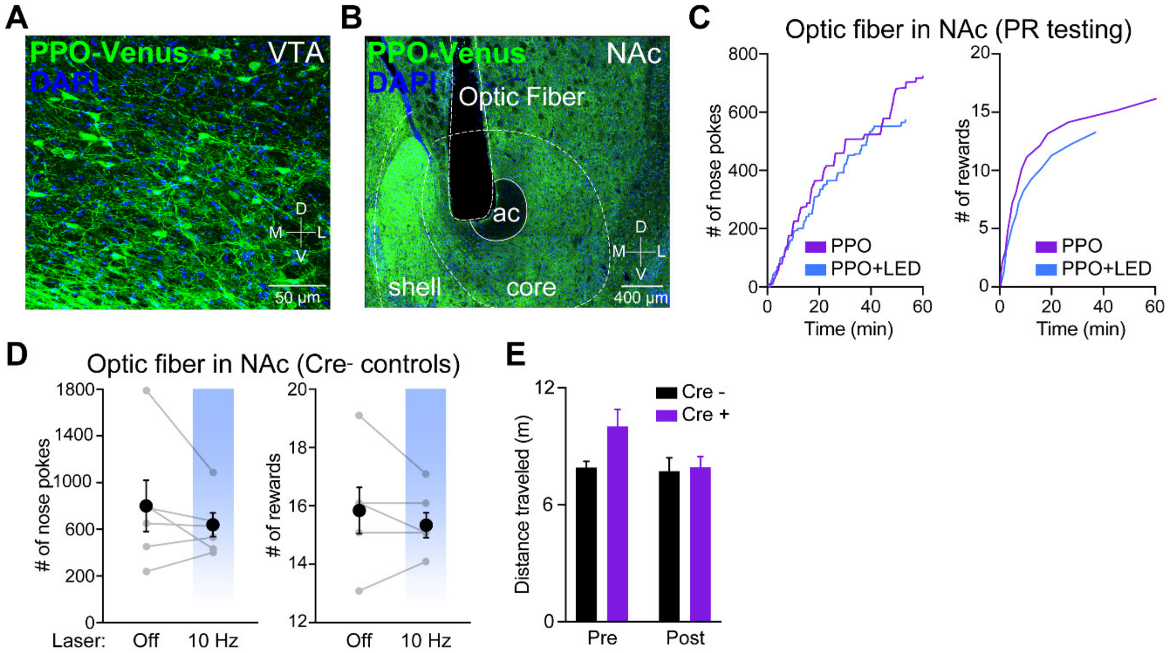


**Supplemental Figure 4. PPO suppresses dopamine neuron-dependent reward behaviors (related to Figure 5).**

(A) Confocal micrograph depicting PPO-Venus expression in DA cell bodies and processes of the VTA in DAT-Cre mice. Scale bar = 50 μm. Dorsal (D), ventral (V), medial (M) and lateral (L) coordinates are indicated.

(B) Confocal micrograph depicting PPO-Venus expression in DA neuron terminals of the NAc. The placement of the optical fiber for targeting these projections is shown. Scale bar = 400 μm. Dorsal (D), ventral (V), medial (M) and lateral (L) coordinates indicated.

(C) Representative time course data for PR nose pokes (left) and rewards obtained (right) over a 60-minute session for one DAT-Cre animal with and without optical stimulation of DA terminals in the NAc

(D) Summary graphs of PR testing in Cre negative animals with and without optical stimulation of DA terminals in the NAc (blue bars). Terminal inhibition decreased the number of rewards that mice received (right) but not the number of nose pokes (left). n=6 mice. p=0.2723

(E) Summary graphs of distance travelled in the pre-conditioning and post-conditioning days while freely exploring the 2 chambers used to test cocaine preference in Cre negative and PPO expressing mice. Cre negative and PPO-expressing mice show no differences in locomotor activity. n=6 for Cre negative and n=5 for PPO-expressing mice. Cre negative vs. PPO-expressing, P=0.06 for pre-conditioning day and p=0.9686, on the post-conditioning day.

**Video S1. PPO is activated by UV and blue light (related to Figure 1).**

Movie of $\gamma$9-mCherry translocation in HeLa cells co-expressing untagged parapinopsin (PPO). $\gamma$9-mCherry (red) was imaged with a 595 nm laser which did not cause spontaneous translocation. Widefield stimulation with blue (470 nm) and UV (365 nm) light (150 ms pulses every 5 seconds was delivered when indicated. Note the rapid translocation of $\gamma$9-mCherry from the plasma membrane to intracellular compartments during photostimulation and recovery when the light is switched off.

**Video S2. PPO can be multiplexed with red fluorescent $Ca^{2+}$ sensors for simultaneous imaging and manipulation of activity (related to Figure 1).**

Movie of fluorescence transients from the red fluorescent $Ca^{2+}$ indicator RCaMP1.07 in cultured striatal neurons co-transfected with PPO-Venus (not shown). Neurons exhibited baseline $Ca^{2+}$ fluorescence transients suggesting spontaneous firing in these cultures and were unaffected by imaging with a 561 nm laser. Blue LED stimulation was delivered in the boxed region of interest

(10 Hz, 10 ms, 10 mW/mm$^2$) which suppressed these transients in the targeted cell and led to increased transient frequency in nearby neurons.

**Video S3. PPO can be activated at single-cell resolution (related to Figure 1H, I).**

Movie of $\gamma$9-mCherry (red) translocation in HeLa cells co-expressing untagged parapinopsin (PPO). LED stimulation of individual cells in the boxed regions of interest was delivered through a digital mirror device. Note the rapid translocation in the regions of interest which did not affect translocation in adjacent cells.
